# Quantifying landscape-flux via single-cell transcriptomics uncovers the underlying mechanism of cell cycle

**DOI:** 10.1101/2023.08.01.551525

**Authors:** Ligang Zhu, Jin Wang

**Affiliations:** College of Physics, Jilin University, Changchun 130021, China; State Key Laboratory of Electroanalytical Chemistry, Changchun Institute of Applied Chemistry, Chinese Academy of Sciences, Changchun 130022, China; Department of Chemistry, Physics and Astronomy, Stony Brook University, Stony Brook, NY 11794, USA

## Abstract

Recent developments in single-cell sequencing technology enable the acquisition of the whole transcriptome data. However, understanding the underlying mechanism and identifying the driving force of the transcriptional regulation of the cell function directly from these data remains challenging. To address this urgent need, we reconstruct a continuous vector field of cell cycle based on the discrete single-cell RNA velocity to quantify the single-cell global non-equilibrium dynamic landscape-flux. We reveal that large fluctuations disrupt the global landscape and genetic perturbations alter landscape-flux, thus identifying key genes in maintaining cell cycle dynamics and predicting associated effects on function. Additionally, we quantify the fundamental energy cost of the cell cycle initiation and reveal that sustaining the cell cycle requires curl flux and dissipation to maintain the oscillatory phase coherence. We enable the inference of the cell cycle gene regulatory networks directly from the single-cell transcriptomic data, including the feedback mechanisms and interaction intensity. This provides a golden opportunity to experimentally verify the landscape-flux theory and also obtain its associated quantifications. Our study also offers a unique framework for combining the landscape-flux theory and single-cell high-through sequencing experiments together for understanding the underlying mechanisms of the cell cycle and can be extended to other non-equilibrium biological processes, such as differentiation-development and disease pathogenesis.

## Introduction

The cell cycle is a complex process that controls cellular growth and division, playing a crucial role in the development and maintenance of living organisms (1, 2). An improved understanding of the regulation of the cell cycle is essential for comprehending various biological processes, including cellular differentiation, tissue development, and disease pathogenesis. Integrating statistical mechanics and single-cell biology has provided novel opportunities to study the cell cycle and its regulation (3). Statistical mechanics offers a powerful framework for investigating complex systems, molecular interactions, and driving forces underlying cellular processes (4). Contrarily, single-cell biology provides a method to explore cellular processes at the single-cell level, which facilitates the comprehension of the heterogeneity of the cell populations and the mechanisms of cell differentiation (5). The precise study of the cell cycle is vital for understanding the life of a single cell, which is the fundamental unit of living systems. Conventionally, researchers explore the dynamics of the trajectories or perform local stability analysis around fixed points for the nonlinear system. However, this does not give a global picture or global stability, so people search further for landscapes to characterize the global stability of the system. However, the landscape alone can only quantify the dynamics of the equilibrium system with detailed balance (no net input to or output from), but it cannot describe the whole dynamics of the non-equilibrium system, an additional rotational flux as the driving force is necessary as we pointed out (6). The transitions between the gene expression states, which determine the driving forces, arise from a non-equilibrium effective potential landscape caused by the steady-state probability and the rotational steady-state probability flux between system states (7). The steady-state probability flux measures the degree of the non-equilibrium and the irreversible behavior of a system, linking the dynamical and thermodynamic aspects of the system. Since flux originates from energy input and generates thermodynamic dissipation, it can act as a dynamic driving force (8). However, prior studies modeled the dynamics of cellular signaling networks based on the existing knowledge of gene regulatory networks, necessitating the estimation of several model parameters and network model simplification, deviating significantly from the real biological systems (9, 10). With advancements in single-cell sequencing technology, complete transcriptomic data can be acquired. Several studies constructed the cell fate landscape from the state manifold of the single cell, which did not obtain the underlying mechanism of the cell signaling regulation (3, 11–16). Consequently, studying the underlying mechanism via non-equilibrium dynamics and thermodynamics of transcriptional regulation of cell function directly from single-cell transcriptomic data demands urgent attention.

Our work aims to contribute to the understanding of the cell cycle mechanism via nonequilibrium dynamics and thermodynamics by investigating the landscape-flux quantified from the single-cell transcriptomic data. Single-cell RNA sequencing (scRNA-seq) provides information on the gene expressions in the individual cells and enables us to analyze the gene expressions during the cell cycle. Through assessing the single-cell RNA velocity, reconstructing vector fields, reducing dimensions, mapping these patterns to lower-dimensional spaces, we can visualize the landscape-flux and understand the complex interactions between the different signaling processes that regulate the cell cycle, as well as the dynamic flux and thermodynamic dissipation that drive and maintain the cell cycle processes. We then performed in silico perturbations that enable us to identify key genes for cell cycle regulation and predict the effects of their genetic perturbations on the cell cycle dynamics and thermodynamics. Calculating the phase diffusion, coherence, and coherence time of the cell cycle oscillations, we quantified their relationships with dynamic flux and thermodynamic consumption. We found that noise mediates phase diffusion and reduces coherence and coherence time, resulting in shorter or even impossible cell cycle maintenance times. However, living cells continuously fight to maintain cell cycle oscillation by consuming energy, leading to a non-equilibrium process where entropy increases. We also found that the dynamic flux plays a role in maintaining the continuous cell cycle, which dynamically drives the cell continuously into the next cell cycle phase and promotes the coherent oscillation (cell cycle forward) to maintain the cell cycle oscillation. Lastly, we constructed a transcription factor regulatory network by identifying differentially-expressed genes between different cell cycle phases and validated the key gene circuit behind cell cycle oscillation dynamics by calculating the Jacobian.

The cell cycle is crucial for cell replication and division, governing cellular proliferation and development, which are the foundation of life. Our study employs a unique combination of experimental high-throughput sequencing and theoretical landscape-flux approaches to understand the origin and driving forces of the cell cycle. The purpose of our research is to utilize high-throughput data to experimentally verify the landscape and flux theory and provides its associated quantifications. Computational analysis identifies key genes and regulators critical for normal cellular function and compares them with the experimental data. Our work enhances our understanding of the biological functions of the cell cycle and has implications for maintaining normal cell function, which is crucial in disease prevention and human health. Moreover, the global principles from non-equilibrium dynamics and thermodynamics can be used to pin down the mechanisms in regulating cell cycle processes across organisms and underlying regulatory networks. In summary, our research represents an essential step forward towards comprehending the cell cycle and its regulation, potentially having a significant impact on the field of single-cell biology and beyond.

## Results

### Quantifying cell cycle non-equilibrium landscape-flux from single-cell transcriptomics data

Traditional systems biology studies have relied on a priori knowledge of biological regulatory networks to model and study cell dynamics; however, this approach is associated with several limitations. One major limitation lies in the simplification of networks, which may lead to an incomplete understanding of complex biological systems (17). Additionally, obtaining reliable results from these models can be challenging due to the estimation of a large number of parameters, such that they may not accurately reflect the real biological system (10). Although some studies have constructed a cell fate landscape from the state manifold of single cells, they are unable to obtain the underlying physical mechanism related to cell signaling regulation (13–16). To overcome these limitations, we present a novel approach to studying the underlying mechanism of the cell cycle via non-equilibrium dynamics and thermodynamics by bypassing the prior known regulatory networks and quantifying the cell cycle regulatory dynamics directly from the experimental data collected by single-cell high throughput sequencing.

We quantified the cell cycle dynamics landscape-flux from the single-cell transcriptomics data, following four primary steps (Figure 1A). Firstly, we downloaded and processed the cell cycle transcriptomics data, which provided information on gene expression patterns of individual cells over time for subsequent analysis. Secondly, we estimated RNA velocity from transcriptomics data. Thirdly, we reconstructed the vector field of cell cycle dynamics from the RNA velocity information to visualize the movements of cells throughout the cell cycle. This captured the overall flow of cells through the cell cycle and provided us with a way to visualize the movements of cells through the cell cycle. In the final step, the landscape-flux of global cell cycle dynamics and thermodynamics was quantified. This method included UMAP-based identification of cell cycle phase clusters, calculation of RNA velocity, reconstruction of cell cycle dynamics’ vector field, computation of the cell cycle dynamic potential landscape, curl flux of the cell cycle dynamic landscape, and gradient force assessment of the cell cycle dynamic landscape.

**Figure 1.**
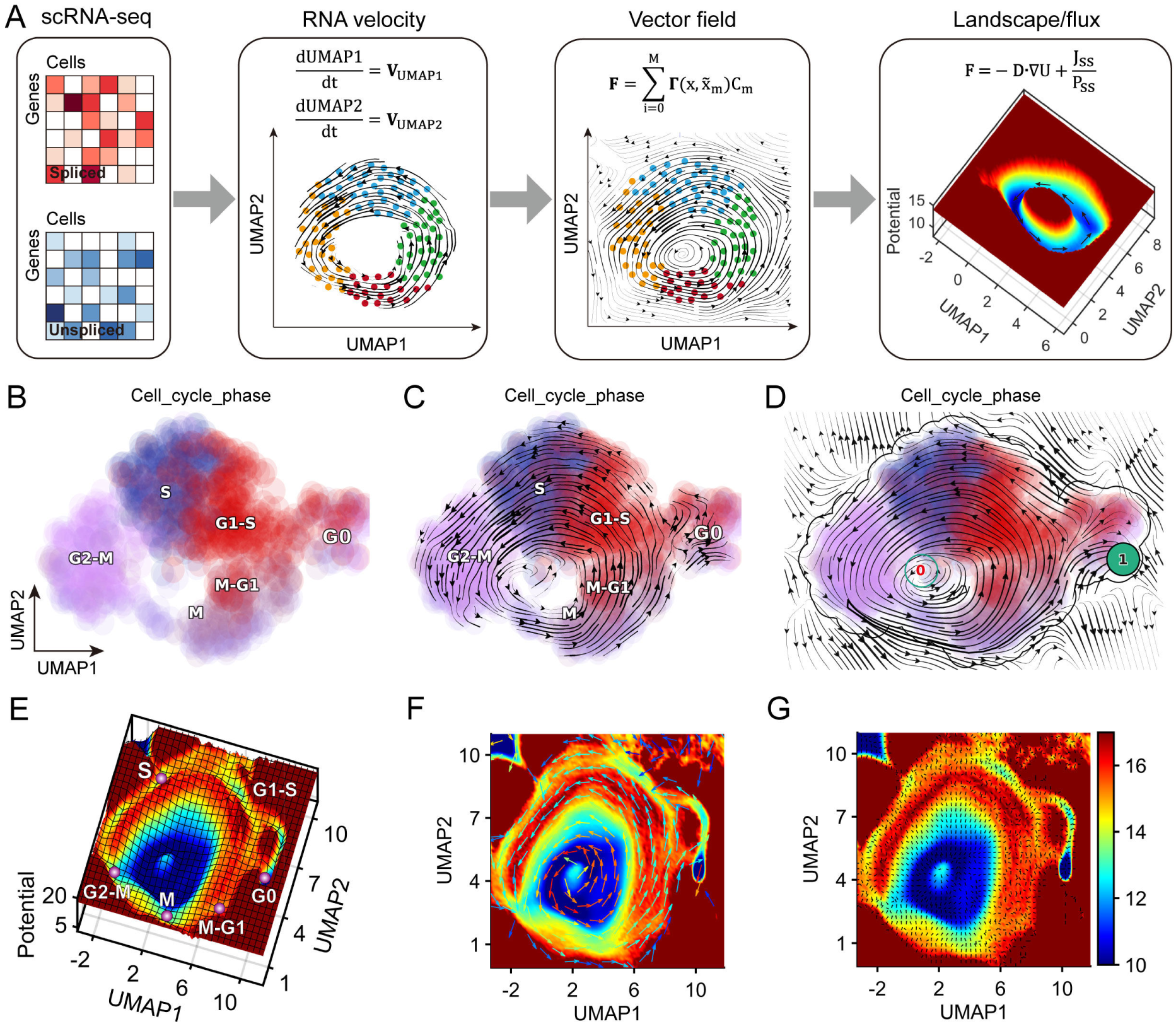
Workflow for analyzing cell cycle landscape-flux from single-cell transcriptomics data. (A) Workflow of constructing cell cycle landscape-flux by single-cell transcriptomics data. (B) Cell cycle phase clusters by UMAP of U2OS cells. (C) RNA velocity of cell cycle dynamics of U2OS cells. (D) Reconstructed vector field of cell cycle dynamics of U2OS cells. (E) Potential landscape of cell cycle dynamics in UMAP of U2OS cells. (F) Curl flux (colored arrow) of cell cycle dynamics landscape in UMAP of U2OS cells. The color of the arrow presents the magnitude of flux, red corresponds to larger, and blue to smaller. (G) The gradient force (black arrow) of cell cycle dynamics landscape in UMAP of U2OS cells.

To gain a deeper understanding of the underlying structure and flow of cells through the cell cycle, we performed a landscape-flux analysis of the cell cycle dynamics using scRNA-seq data from human bone osteosarcoma epithelial cells (U2OS cells)(18) and single-cell 5-ethynyl-uridine-labeled RNA sequencing (scEU-seq) data from human retinal pigment epithelial-1 cells (RPE1 cells)(19). We first identified cell cycle phase clusters in U2OS cells by applying the UMAP algorithm (20) to generate a low-dimensional representation of the cells (Figure 1B). Cells were then colored based on their cell cycle phase, allowing us to visualize the distribution of the cells in the cell cycle and to observe the heterogeneity of the cell cycle phases. The expressions of the transcription factors in the cells are different in different cell cycle phases, and the gene regulations are also different, so the cells have different cycle states (Figure S1). RNA velocity (21, 22) was then calculated for each cell to capture the direction and magnitude of changes in the gene expressions (Figure 1C), which facilitated visualization of phase portraits and insight into cell cycle progression. The vector field of cell cycle dynamics was reconstructed using Dynamo (23), which allows us to reconstruct the function of the continuous and analytic velocity vector field from sparse, noisy single-cell velocity measurements (Figure 1D). At the same time, we performed a differential geometry analysis of the cell cycle based on this vector field (24). Speed defines the length of the velocity vector in the vector field, that is, the magnitude of the velocity, which represents the magnitude of the force in the Langevin equation (Figure S2B). Divergence characterizes the local flux exiting versus entering an infinitesimal region in the expression space - the “outgoingness”. The sources (sinks) often have strong positive (negative) divergence. The divergence in the cell cycle vector field represents the degree of convergence, reflecting the distribution of different cell cycle phases in the vector field (Figure S2C). Curl characterizes the infinitesimal rotation of a cell state in the vector field, which is gradient-free and represents the cell cycle state. The U2OS cell cycle velocity vector field is noise-free, so the limit cycle of the cell cycle oscillation dynamics is small, and the curl is larger at the small limit cycle (Figure S2D). The acceleration field reveals hotspots of cell states where the velocities change dramatically. When a cell leaves one cycle phase and enters another cycle phase, the acceleration is high. Acceleration represents an early driver of cell cycle regulation (Figure S2E). The curvature field reveals hotspots where the velocity abruptly changes direction, such as regions around unstable fixed points where the gene expression changes from activation to repression or vice versa. Genes that contribute strongly to the curvature are drivers that control the cell fate. The curvature reflects the cell cycle phase bifurcation, which is larger in the G0 phase, indicating the switch between the oscillation-limited cycle of the maintenance cycle and the G0 monostable state of the arrest cell cycle (Figure S2F). Collectively, these results complementary revealed hotspots of the cell states where velocities change dramatically, highlighting key transition points and driving forces of the cell cycle, including stability and instability. Using the reconstructed function, we also quantified the potential landscape U (U = −ln *P_ss_*) of the global cell cycle dynamics in UMAP, representing the steady state probability distribution *P_ss_* of cells at different locations in different cell cycle phases with the shape of an irregular Mexican hat (Figure 1E). The Mexican hat-shaped landscape features two ring valleys, and the small ring valley in the inner blue area is a virtual loop, and the real cell cycle oscillation trajectory corresponds to the outer ring valley (25). The emergence of such two different limit sets is a common phenomenon when performing dimensionality reduction with limit cycles to UMAP plane, which is reminiscent of the Poincaŕe-Bendixson theorem in planar dynamics theory (26). The different phases of the cell cycle correspond to different basins of attractions (Purple balls) along the outer close oscillation ring valley path in the Mexcian landscape. Inside or outside the Mexican hat outer ring, the potential is higher and the probability of the system reaching these areas is low. The basin depth quantifies the duration of the stay in the corresponding phases and the local barrier between these phases or basins corresponds to the check point of the cell cycle (9). While the landscape attracts the system down to the cell cycle ring valley from the gradient part of the driving force, it is the curl flux from the rotational part of the driving force which drives the coherent oscillation of the cell cycle. Therefore, the Mexican hat shape landscape and the curl flux together guarantee the stability and the robustness of cell cycle oscillation dynamics. This potential landscape also reflects the stability of different states in the cell cycle and highlights key transition points. Additionally, we quantified the curl flux and negative gradient force of the cell cycle nonequilibrium dynamics landscape in UMAP from the reconstructed function (Figure 1F and 1G). The curl flux indicates the amount of rotational flow in the cell cycle dynamics landscape that drives the cell cycle oscillations along the closed ring valley path, while the negative gradient force indicates the direction and magnitude of the forces stabilizing the cell cycle phase. If the driving force from the flux is greater than the force from the negative potential gradient, the cell will go across the G0 basin barrier into the G1-S phase and the cell will enter the cycle phase. Otherwise, the cell will return to the G0 basin and arrest in the G0 state. Biologically, this means that the nutrient supply is not large enough, or the DNA is damaged, as in the case of senescent cells (27).

Similarly, we evaluated the total RNA velocity from the scEU-seq of the RPE1 cell cycle, reconstructed the vector field for further differential geometric analysis (Figure S4), and quantified the non-equilibrium landscape-flux (Figure S3). These findings verify the presence of the curl flux as another component of the driving force for the nonequilibrium dynamics in additional to the gradient of the cell cycle landscape by experimental single-cell high throughput sequencing data, providing deeper insights into the mechanisms behind the stability and behavior of intrinsic nonequilibrium cell cycle progression that cannot be discerned by the landscape quantification alone. In fact, the cell cycle is one of the best examples of demonstrating the nonequilibrium dynamics being dictated by both the landscape and the flux. What’s more, the creation of the cell cycle non-equilibrium landscape-flux through single-cell transcriptomics data presents a new and powerful tool for the study of cell cycle dynamics, allowing for insights into underlying mechanisms that drive cell cycle progression.

### Noise alters the global cell cycle dynamics and thermodynamics

Variability in biomolecular noise or changes in molecular states can affect intracellular gene regulations and networks, which are crucial for the uncertain, multi-destiny behavior of the system (28–30). To investigate the effects of noise intensity on the underlying mechanism of the cell cycle via nonequilibrium dynamics and thermodynamics, we performed simulations at varying noise intensities and analyzed the resulting global changes in the cell cycle dynamics and thermodynamics. Firstly, we examined changes to the global cell cycle dynamic landscape and show that as the noise intensity increases, the global dynamics landscape of the cell cycle becomes more complex and less predictable. Specifically, with zero noise, the cell cycle progresses very smoothly and predictably (Figure 2A), while the ring valley of cell cycle becomes wider as noise intensity increases (Figure 2B and 2C), whereas the ring valley is broken when the noise is high, showing the discrete attraction basins, at which time the cell cycle is broken and the cells arrest in the different phases (Figure 2D). Moreover, we found that the barrier height decreased in the G0 phase and oscillation center in the landscape as noise intensity increased (Figure 2I). Barrier height of G0 phase is defined as the barrier from the G0 basin to the saddle point (between G0 and G1-S), which quantifies how easily cells can switch from quiescent G0 phase to the cyclic state. The barrier height of the oscillation center is defined as the potential difference between the minimum potential of the landscape and the local maximum inside the limit cycle, which characterizes the global stability of the oscillation system (7). We also investigated and found that the noise increases the cell cycle period, indicative of the rate at which the cell cycle oscillates and how fast it grows (Figures 2J and S6). Oscillation amplitude of UMAP1 increases with the noise intensity, resulting in a wider range of the oscillation limit cycle (Figures 2J, S5 and S6). Increased variability or noise effectively makes potential landscape basins shallower, thereby broadening the distribution of biological variables (31). By quantifying the trajectory autocorrelation function and the corresponding power spectrum density of the oscillating trajectory for different diffusion coefficient D, we can see that as the noise increases (the stability of the oscillation system decreases), the peak height of the power spectrum density decreases and the distribution of the power spectrum becomes more spread out (Figure S5). This indicates that the peak height and the width of the power spectrum density can be a measure of the stability of the oscillation system. To be clear, when D is in the range of 0.01 to 0.1, the landscape is more realistic, with obvious cell cycle checkpoints, and it is consistent with the true level of experimental measurements (32, 33).

**Figure 2.**
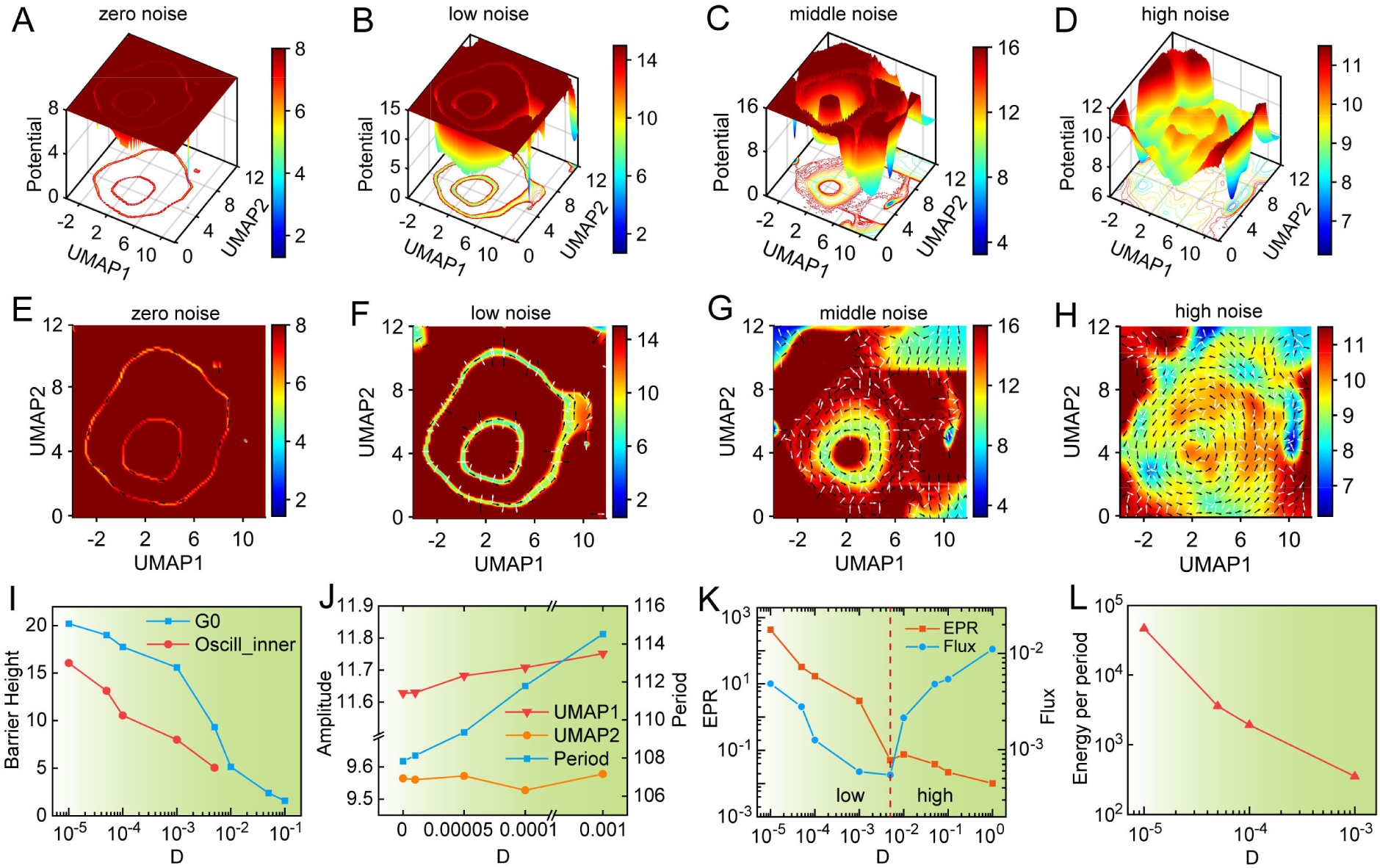
Cell cycle nonequilibrium landscape-flux affected by noise intensity. (A to D) Cell cycle global dynamics landscape of U2OS cells changes in UMAP when noise intensity is 0 (A), 0.00005 (B), 0.005 (C), 0.05 (D). (E to H) Curl flux (black arrow) of cell cycle dynamics landscape in UMAP when noise intensity is 0 (E), 0.00005 (F), 0.005 (G), 0.05 (H). (I) The change of Barrier Height of G0 phase (blue) and oscillation center (red) in landscape when the noise intensity is changed. (J) The change of amplitude of UMAP1 and UMAP1 and period of cell cycle oscillatory dynamics when the noise intensity is changed. (K) The change of EPR and Flux of cell cycle when the noise intensity is changed. (L) The change of Energy per period of cell cycle when the noise intensity is changed.

Furthermore, we quantified the curl flux as a major component of the driving force of the cell cycle and the entropy production rate (EPR) as the measure of the thermodynamic cost association to the nonequilibrium curl flux of the cell cycle dynamic landscape to investigate how noise intensity affected the underlying mechanism of the cell cycle via nonequilibrium dynamics and thermodynamics. As shown in Figure 2E-H, increasing noise intensity resulted in a more widely distributed flux, with cells flowing between different phases less predictably. We observed that in the small noise range, the flux decreases as the noise increases, while the cell cycle is still oscillating (Figure 2K). Larger flux derived from the energy input or nutrient supply implies a larger cell cycle driving force and subsequently results in a shorter time to complete the cell cycle oscillation (period is smaller). Indeed, the speed of the cell cycle is a hallmark of cancer. However, when the noise is very large, although the flux becomes larger with the increase of noise, the cell cycle has been destroyed. At this point, the flux drive on the cell cycle can no longer resist the destruction of noise. In addition to global dynamics landscape alteration, noise intensity also affects the thermodynamics and energetics of the cell cycle. We found that the EPR of the cell cycle decreases with increasing noise intensity (Figure 2K). Furthermore, the energy per period of the cell cycle decreases with increasing noise intensity (Figure 2L), indicating a change in the energy requirements of the system. These results imply that the cell cycle becomes less efficient at higher noise levels, suggesting that the system requires energy to maintain the coherent cell cycle oscillations.

Taken together, these results demonstrate that noise intensity alters the underlying mechanism of the cell cycle via global nonequilibrium dynamics and thermodynamics, suggesting that the feedback of intracellular gene interactions plays an important role in maintaining the robustness and stability of the cell cycle in the face of environmental fluctuations. These findings have crucial implications for our understanding of the mechanisms that control cell cycle progression and may have important applications in the development of novel therapeutic strategies for the treatment of diseases associated with cell cycle dysregulation. These changes suggest that noise intensity may play a critical role in regulating the onset and termination of cell cycle phases.

### *In silico* perturbation analysis cell cycle dynamics and thermodynamics under genetic perturbation

Genetic perturbation refers to the alteration of the function of a biological system by external or internal factors, such as environmental stimuli, drug inhibition, and gene knockdown (34). It is used to study how genes interact with each other and affect cellular processes and phenotypes (35–37). To investigate the effects of the genetic perturbations on the underlying mechanism of the cell cycle via nonequilibrium dynamics and thermodynamics, we performed *in silico* perturbations of different genes in a controlled and systematic manner. Key genes involved in cell cycle regulation, such as cyclin-dependent kinases (CDKs), cyclins, and tumor suppressor genes, were perturbed in this study. Similar to Figure 1, we first estimated the RNA velocity of U2OS cells under different genetic backgrounds by *in silico* perturbation (Figure S7A), and then reconstructed the different vector fields based on the perturbed RNA velocity (Figure S7B). Finally, we quantified the global cell cycle dynamic landscapes under different genetic perturbations by adding noise in the dynamic simulation (Figure 3A). We found that perturbing distinct genes resulted in specific changes in the cell cycle landscape and alterations in EPR and flux (Figure 3B and 3C).

**Figure 3.**
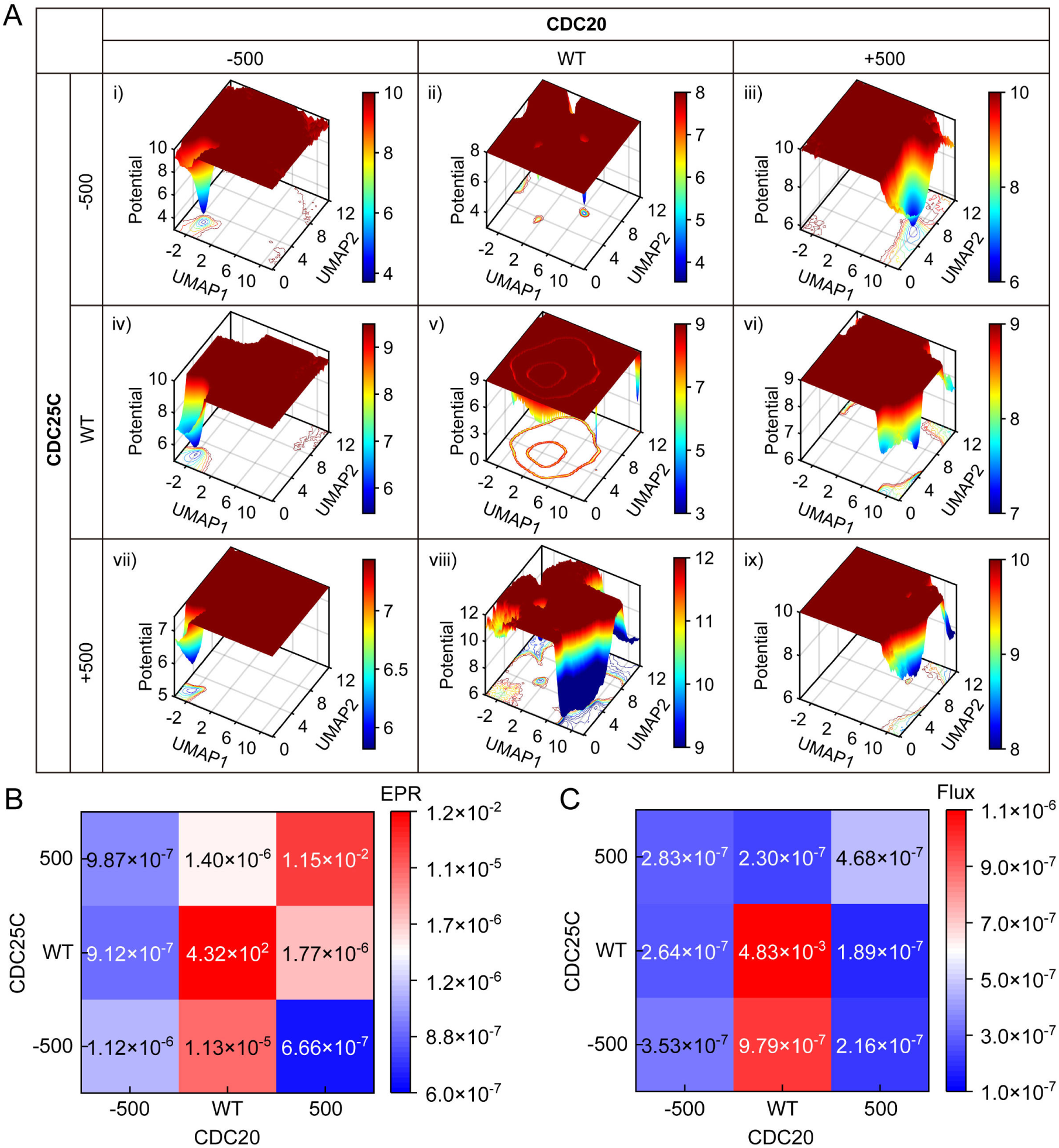
The genetic perturbation alters cell cycle landscape/flux. (A) Cell cycle global dynamics landscape of U2OS cells in UMAP with different genetic perturbation. Validation of in silico trajectory predictions. (i) Suppression of both CDC20 and CDC25C. (ii) Suppression of CDC25C only. (iii) Activation of CDC20 and suppression of CDC25C. (iv) Suppression of CDC20 only. (v) WT. (vi) Activation of CDC20 only. (vii) Suppression of CDC20 and activation of CDC25C. (viii) Activation of CDC25C only. (ix) Activation of both CDC20 and CDC25C. (B) EPR of cell cycle nonequilibrium thermodynamics with different genetic perturbation. (C) The average Flux of cell cycle nonequilibrium dynamics with different genetic perturbation.

Specifically, the negative perturbation of *CDC20* led to an increase in the barrier height of the G2-M phase with higher chance of arrest in this phase, while positive perturbation of *CDC20* led to a deeper basin of the G1 phase with higher chance of arrest in this phase (Figure 3A). As the experiment showed, *CDC20* plays a crucial role in cell division, being the target protein in the spindle checkpoint during mitosis (38). We also found that the negative perturbation of *CDC25C* caused disruption of cell cycle oscillation and led to a deeper well of the G2-M phase leading to possible arrest, albeit less obviously than the effect caused by perturbing *CDC20* (Figure 3A). Biologically, the downregulation of *CDC25C* induces cell cycle arrest in G2/M phase in response to DNA damage via p53-mediated signal transduction, and its abnormal expression is associated with cancer initiation, development, metastasis, occurrence and poor prognosis (39). Similarly, genetic perturbations in RPE1 cells are manifested that the negative perturbation of *KPNA2* led to a higher barrier height of the G2-M phase, whereas the positive perturbation of *CDC20* led to a higher barrier height of the M phase (Figure S8A). This finding aligns with experimental evidence: *KPNA2* knockdown downregulated *CCNB2* and *CDK1*, inhibited cell proliferation, and induced cell cycle arrest in the G2-M phase (40). *CCNE2* is a regulatory subunit of cyclin-dependent kinase 2 (Cdk2) and is thought to control the transition of quiescent cells into the cell cycle (41). Our genetic perturbations also confirm the cell cycle initiation function of *CCNE2*, as reflected in the deepening of the G0 well under negative perturbation of *CCNE2*, and disappearance of the G0 basin with the appearance of other periodic phases following positive perturbation of *CCNE2* (Figure S8A).

Figure 3B shows the EPR of cell cycle nonequilibrium thermodynamics with different genetic perturbations. We observed that different genetic perturbations can cause significant changes in EPR, with some resulting in increased energy consumption and others in decreased consumption. However, the EPR is lower under genetic perturbations than in WT, strong perturbations disrupt cell oscillatory dynamics (Figures 3B and S8B). Thus, maintaining continuous cell oscillation requires greater energetic costs than arresting in different periodic phases. We also quantified the average flux of cell cycle nonequilibrium dynamics with various genetic perturbations and found that WT cells exhibit the largest curl flux, where the flux as a driving force of the cell cycle is greater than the negative gradient force, and the flux drives the cell cycle to continue moving forward to maintain the coherent oscillation (Figures 3C and S8C). In contrast, specific genetic perturbations result in a flux less than the negative gradient force, insufficient to drive the cell cycle forward, and causing cells to arrest at certain cell cycle phases.

Overall, the results demonstrate the efficacy of using the reconstructed vector field based on RNA velocity derived from single-cell omics data of the cell cycle to predict how genetic perturbations impact cell cycle behavior. This methodology has the potential for identifying the critical regulatory genes involved in maintaining cell cycle coherent oscillation, such as *CDC25C*, *CCNE2*, *KPNA2*, etc., which may be the new targets in cancer.

### The underlying mechanism of cell cycle initiation and termination via non-equilibrium dynamics and thermodynamics

The ability to drive cellular state transitions has gained attention as a promising strategy for disease modelling (42). The quantified Waddington landscape provides a way of describing and revealing biological pathways of developmental processes (43). A previous study confirmed that gradient and curl forces control the dynamics of these biological paths on the landscape, rather than following the steepest descent path expected (17). To investigate the cell cycle progression and transitions between different phases, we computed the Least Action Paths (LAPs) and Mean First Passage Time (MFPT) between different cell cycle phases. LAPs are optimal phase space trajectories minimizing the action’s cost of moving from one state to another, while MFPT is the average transition time between states. Specifically, the optimal path between any two cell states (e.g., the fixed points of G0 phase and G1/S phase) is searched by changing the continuous path connecting the source and target states while minimizing its effect and updating the associated transition time (9). The resulting LAP has the highest transition probability and is correlated with a specific transition time.

We first map the LAPs between different cell cycle phases in the velocity vector field, where the nodes along the paths represent the discrete points, with color indicating LAP transition time and arrows depicting path direction, and the black curves connecting different phases represent LAPs (Figures 4A and S9D). We also calculated and illustrated the LAPs on the landscape, which reveals the optimal path (least action) that cells take to overcome potential barriers during the cell cycle (Figures 4B and S9E). Green curves connect G0 to G1-S and M to G0, representing the cell cycle’s initiation and termination, respectively. The cell cycle path is non-uniform and sandwiched between flux (black arrows) and negative potential gradient (white arrows), demonstrating that progression arises from both curl flux and negative potential gradient force. To enter the cycle from the G0 basin to the G1-S basin, the cells must overcome a barrier on the path (saddle point on the landscape) located between the G0 and G1-S basins, where the flux acts as a driving force.

**Figure 4.**
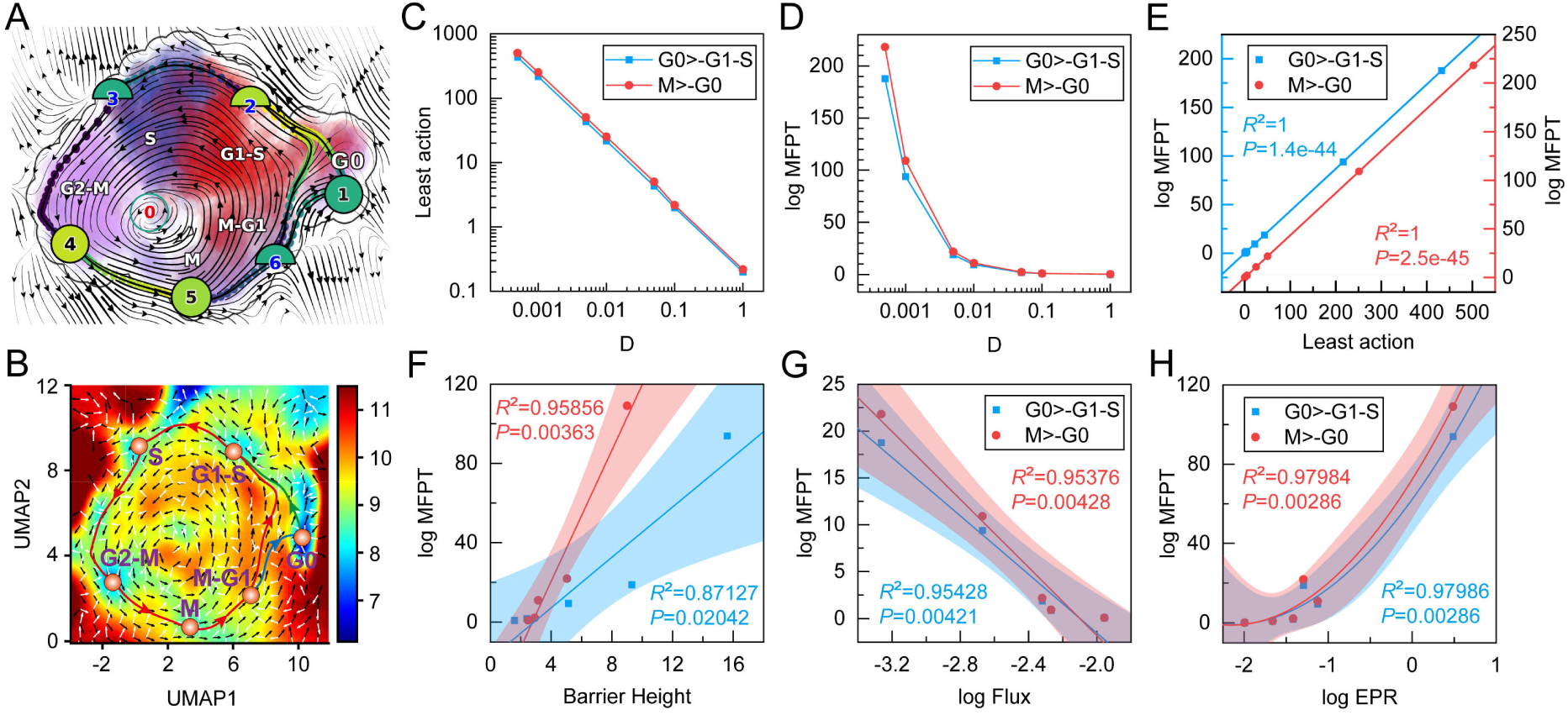
LAPs and MFPT in cell cycle initiation and termination. (A) LAPs between different cell cycle phases in the velocity vector field of U2OS cells. The color of digits in each node reflects the type of fixed point: red, emitting fixed point; black, absorbing fixed point. The color of the numbered nodes corresponds to the confidence of the fixed points. The color of the dots along the paths corresponds to the direction. (B) LAPs between different cell cycle phases in 2D landscape of U2OS cells. The black arrows present the curl flux and the white arrows present the gradient force in the landscape. (C) The least action of G0 phase to G1-S phase and M phase to G0 phase versus diffusion coefficient D. (D) The logarithm of MFPT of G0 phase to G1-S phase and M phase to G0 phase versus diffusion coefficient D. (E) The correlation between the least action and the logarithm of MFPT when diffusion coefficients (fluctuations) are changed. (F) The correlation between barrier height and the logarithm of MFPT when diffusion coefficients are changed. The line is a fitting correlation line, and the shaded part is the 95% fitting confidence interval. (G) The correlation between flux and the logarithm of MFPT when diffusion coefficients are changed. (H) The correlation between the logarithm of EPR and the logarithm of MFPT when diffusion coefficients are changed.

In addition to calculating the LAPs, we also quantified the time of the cell cycle initiation and termination (Figure 4C-4E). We indicated that noise reduces the least actions of cell cycle initiation and termination, resulting in a decrease in MFPT, suggesting that strong fluctuations make it easier for cells to enter the cycle, but also easier to terminate the cell cycle when the cycle coherence is weakened. Furthermore, we found that the logarithm of MFPT is directly proportional to the barrier height on the landscape, i.e. the higher the barrier, the more stable the cell state, and the more difficult it is for the cell to leave this cycle phase (the larger the least action and the longer the escape time) (Figures 4F and S9A). The barrier heights between the attraction basins correlate with the escape time, which characterizes the stability of the cell cycle phase. Additionally, the logarithm of MFPT is negatively correlated with the logarithm of non-equilibrium dynamic flux, suggesting that the larger the flux, the stronger the driving force, and the easier it is for cells to escape a given steady state (Figure 4G). However, thermodynamic energy cost-wise, the logarithm of MFPT is positively correlated with the logarithm of EPR (Figure 4H). The longer the cell takes to escape, the more energy it consumes. To overcome this barrier, cells must expend a certain amount of energy, known as cell cycle initiation energy (44). In summary, these results provide a quantitative assessment of the initial energy required for the progression of the cell cycle (e.g., nutrient supply capacity and metabolic activity (45)) and provide graphical, physical, and quantitative explanations for the checkpoint mechanisms of the cell cycle.

### Maintenance of the cell cycle requires energy-dissipation and flux to enhance the coherence of the oscillation phase

Biological systems need to function accurately in the presence of strong noise, and various regulatory mechanisms have evolved to suppress the fluctuation and environmental effects in processing vital life processes such as cell cycle and development (46). In a noisy environment, cell cycle oscillations can be highly inaccurate due to phase fluctuation, the phase of oscillation follows diffusive dynamics (47, 48). A previous study found that phase coherence is a property of waves that describes how well-aligned the phases of the waves are. When waves are perfectly coherent, their peaks and troughs align precisely, creating a stable and predictable pattern (7, 49). We quantified the relationship between noise intensity and cell cycle oscillation properties (phase diffusion, phase coherence, and coherent time) and found that noise fluctuations increase phase diffusion, decrease phase coherence, and make the coherence time (cell cycle maintenance time) to become lower (Figure S10). In the system of cell cycle regulation, phase diffusion refers to the random fluctuation of the phase of oscillate waves. After certain times, the oscillations can lose the memory or track of the previous dynamics. This makes it harder to maintain the periodicity. The phase coherence is important for maintaining accurate and reliable oscillations in noisy environments. Coherent time refers to the duration over which a system exhibits coherence, or the ability to maintain a stable and predictable pattern (50). In other words, it is the length of time during which oscillating waves in the cell cycle system remain perfectly aligned. There are significant correlations between these factors: increasing phase diffusion leads to a corresponding decrease in phase coherence, and phase diffusion can reduce the coherent time of the cell cycle system, especially in large fluctuation systems (Figure 5A and 5B). Decreased phase coherence leads to decreased coherent time because less well-aligned waves become less stable and predictable over time (Figure 5C).

**Figure 5.**
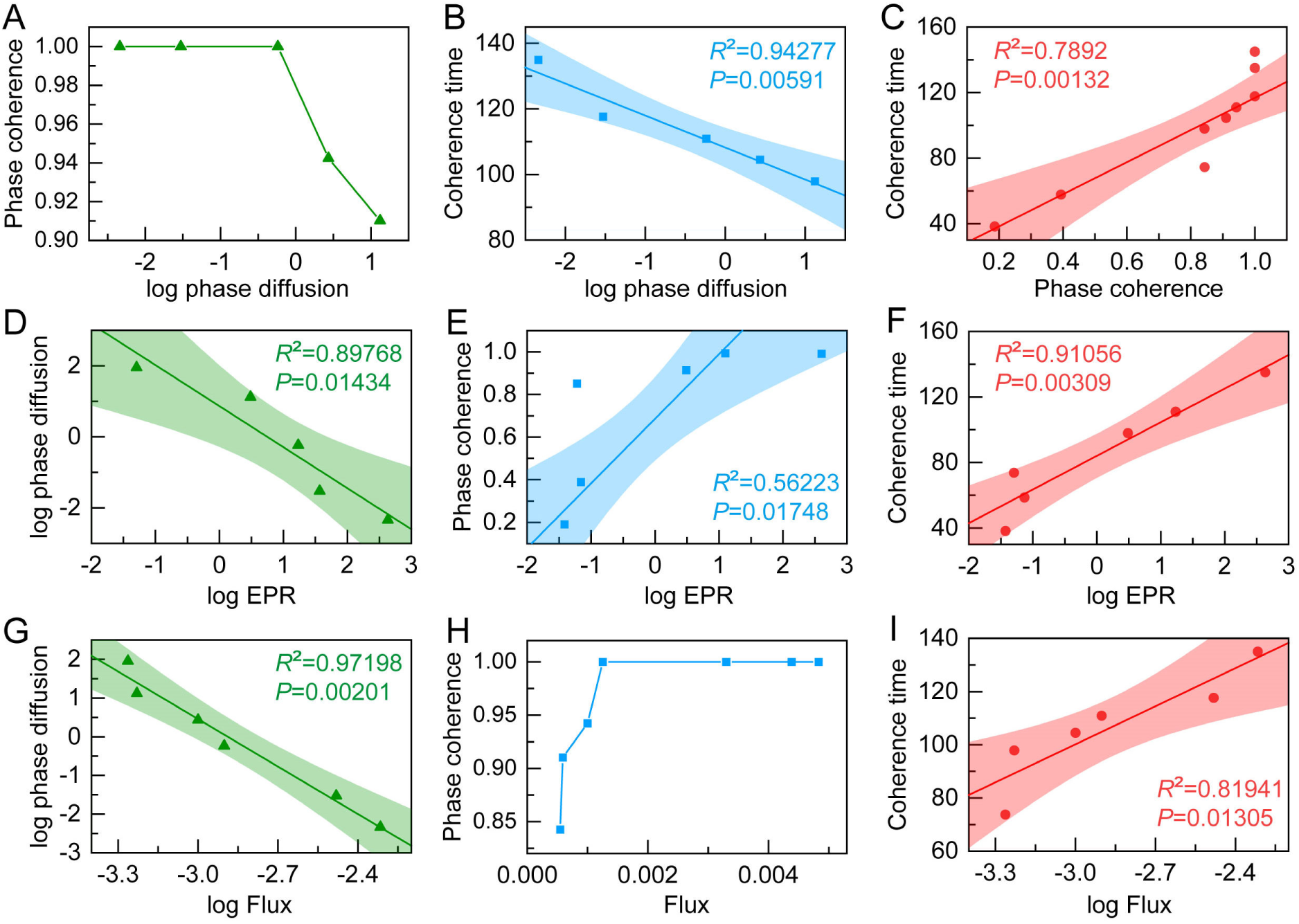
Relationship between phase diffusion, phase coherence, coherence time and EPR and flux. (A) Phase coherence versus the logarithm of phase diffusion. (B) The correlation between coherence time and the logarithm of phase diffusion when diffusion coefficients (fluctuations) are changed. The line is a fitting correlation line, and the shaded part is the 95% fitting confidence interval. (C) The correlation between coherence time and phase coherence when diffusion coefficients are changed. (D) The correlation between the logarithm of phase diffusion and the logarithm of EPR when diffusion coefficients are changed. (E) The correlation between phase coherence and the logarithm of EPR when diffusion coefficients are changed. (F) The correlation between phase coherence and the logarithm of EPR when diffusion coefficients are changed. (G) The correlation between the logarithm of phase diffusion and the logarithm of Flux when diffusion coefficients are changed. (H) The change of phase coherence when Flux is changed. (I) The correlation between coherence time and the logarithm of Flux when diffusion coefficients are changed.

To identify the non-equilibrium mechanism of the cell maintenance cycle progression, we further investigated the effects of dynamic flux and thermodynamic dissipation on the characteristic features of cell cycle oscillations. Our results indicate that the logarithm of phase diffusion is negatively correlated with the logarithm of EPR, while phase coherence and coherent time are positively correlated with the logarithm of EPR (Figure 5D-5F). This suggests that thermodynamic dissipation plays an essential role in regulating the overall dynamics of the cell cycle oscillation. As cells enter the cycle, metabolic processes generate ATP and other energy molecules to provide thermodynamic energy dissipation to maintain the cell cycle oscillations. This added energy can help counteracting the effects of phase diffusion, maintaining coherence, and resulting in more reliable and effective signaling between different cell cycle phases. We also found that changes in the logarithm of phase diffusion were negatively correlated with changes in the logarithm of dynamic flux (Figure 5G). Furthermore, we observed that higher flux levels were associated with increased phase coherence and coherence time (Figure 5H and 5I). These findings highlight the critical role of non-equilibrium dynamic flux force in increasing the stability of the cell cycle flow by enhancing phase coherence and coherent time.

Overall, these results suggest that there is a cost-performance trade-off for noisy biochemical oscillations, where increasing phase coherence comes at a higher thermodynamic cost, which align with the principles of thermodynamic uncertainty relationship (TUR). TUR establishes a trade-off between the dissipation of energy and the precision of observed quantities. In the context of cell cycle oscillating systems, maintaining coherence in the oscillation phases may require the dissipation of energy to counteract the natural tendency of the system to relax towards equilibrium. The coherent oscillation necessitates energy input to sustain the oscillatory behavior and counteract the dissipative processes that tend to disrupt coherence. the TUR can guide the understanding of how energy allocation influences the coherence and precision of oscillation phases. While the energy cost of an oscillating system can counteract the effects of phase diffusion and maintain coherence over longer periods of time, continuous energy input is required to sustain the system’s oscillations, which can be energetically costly in certain circumstances. The energy cost of cell cycle progression is essential for ensuring that cells divide in a controlled and regulated manner, but excessive energy cost or disruptions in energy metabolism can result in cell cycle defects and potentially serious health problems such as cancer. Therefore, regulating energy metabolism and maintaining cell cycle oscillations are critical for preserving the health and proper function of cells and tissues.

### Cell cycle gene regulatory network inference from single-cell transcriptomics data

Gene expression does not appear in isolation, which is influenced by a complex interplay of regulatory interactions with other genes and small molecules. The discovery of these regulatory interactions is the aim of gene regulatory network (GRN) inference methods (51–57). The use of gene regulatory network modeling to analyze biological dynamic processes is an interdisciplinary field that has been widely studied in systems biology. To decode the regulatory mechanism of cell cycle phases, we first identified 27 and 16 differentially expressed genes of U2OS cell and RPE1 cell, separately (Tables S1 and S2), which were used to construct the genetic network that underlies each cell cycle stage. The analytic results demonstrate that the enriched expression of cycle phase-specific genes regulates and maintains cells into different cycle stages via corresponding niche components (Figures 6A and S12A). The nodes in the network represent genes colored according to their associated cell cycle phase. We also identified key genes for each cell cycle phase and constructed a wiring diagram for the cell cycle network model with these key genes (Figures 6B and S12B). The GRN identifies key genes for each cell cycle phase known to play important roles in different cell cycle phases and biological processes such as DNA replication and mitosis. For instance, *CDC6*, *MCM3*, and *CCNE2* are key regulators of G1/S transition and cell cycle initiation that regulate multiple GRNs (CyclinE and CDK2) in the G1-S phases (41, 58). While the S phase is regulated by *MCM6*, *AURKB*, and *CDK1*, which are related to DNA replication and phosphorylate several key proteins to control the transition from G1 to S phase of the cell cycle (59–61). Other important genes identified include *CDC25C*, *CENPE*, and *KIF23*, which regulate the G2/M checkpoint by affecting mitotic cell cycle (62–64). Additionally, the GRN reveals that the M phase is regulated by *CDC20* (38), *PIF1* (65), and *TPX2* (66), *PDE4B* (67), and *KIF20A* (68), highlighting the crucial role of these factors in controlling the M-G1 phase.

**Figure 6.**
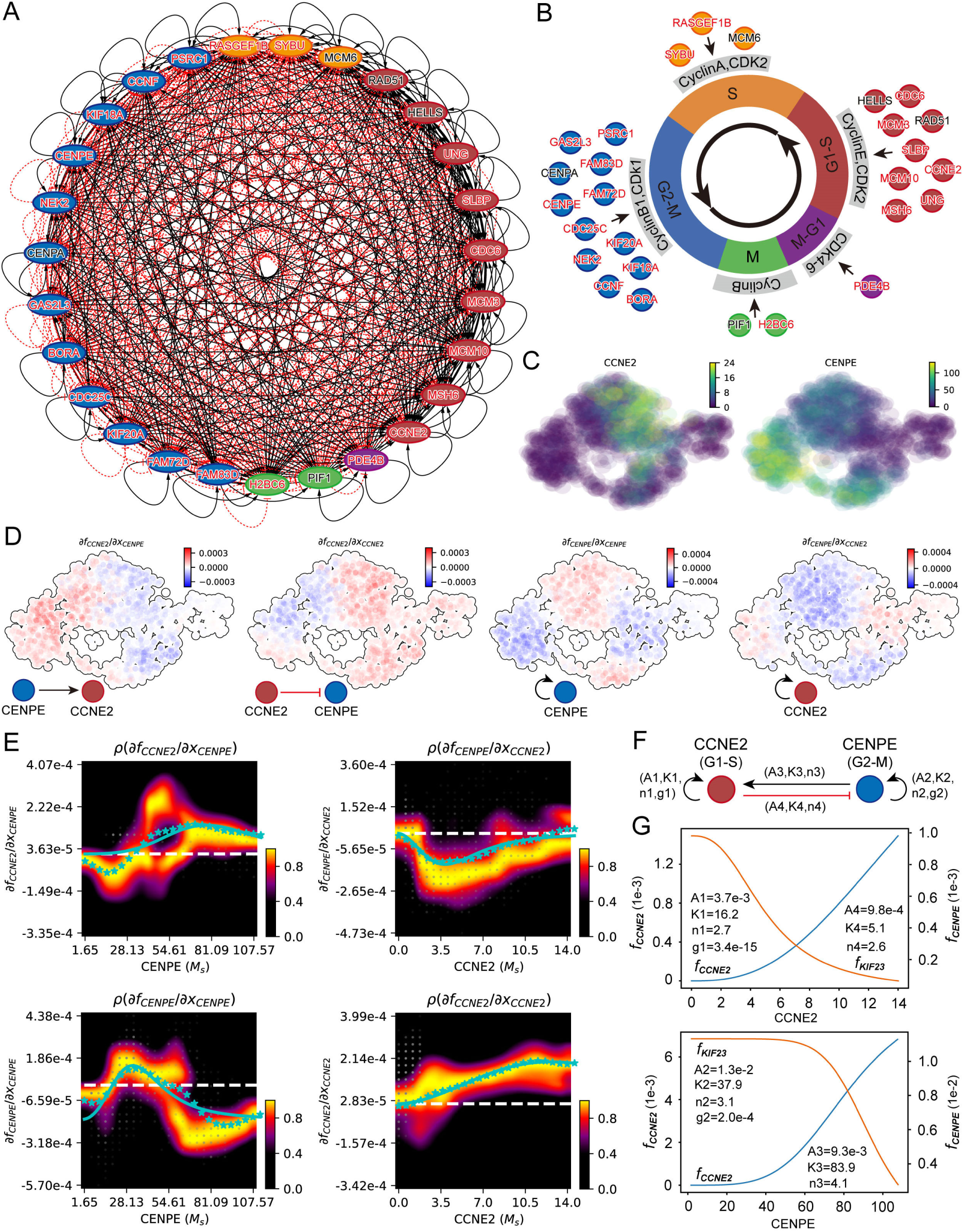
Inference of gene regulatory networks for the cell cycle by single-cell transcriptomics data. (A) The interaction network of genes of each cell cycle phase of U2OS cells. All differentially expressed genes in each cell cycle phase are used to construct network; nodes (genes) with more than one edge are shown. Node colors represent different cycle phases (red: G1-S, yellow: S, blue: G2-M, green: M, purple: M-G1), the black arrows represent the activation and the red arrows represent the inhibition. Genes with black representing TF and red representing non-TFs. (B) The diagram for the cell cycle model with key genes of each cell cycle phase. (C) CCNE2 has high expression in G1-S phase and CENPE has high expression in G2-M phase. (D) Molecular mechanisms underlying the maintenance of the cell cycle. (i) CENPE activates CCNE2. (ii) Repression of CENPE by CCNE2. (iii) Self-activation of CENPE. (iv) Self-activation of CCNE2. (E) Fitting the function of Jacobian versus gene expression with derivatives of a simplistic inhibitory or activation Hill equation. White dashed line corresponds to the zero Jacobian value. The blue stars at each x axis grid point correspond to the weighted mean of the Jacobian values for that point. The blue solid lines are the resultant fittings for the Jacobian. (F) Schematic summarizing the interactions involving CCNE2 and CENPE. (G) The velocity kinetic curves over gene expression changes of the corresponding fitted Hill equations of (E).

Although there are many methods for inferring gene regulatory networks, most of them cannot obtain the strength of regulation, and only a few tools can do it, including Dynamo (23), spliceJAC (53), etc. In our study, we used Dynamo, based on the differential geometry analysis of the reconstructed cell cycle vector field from the single-cell transcriptomics data, we reveal the quantitative information about gene regulation and further uncover the molecular mechanism underlying the maintenance of the cell cycle and the interactions between the key genes. Taking U2OS cells as an example, we demonstrate via our inferred regulatory network that *CCNE2* and *CENPE* regulate the G1-S and G2-M phase, respectively, showing peak expression in their respective regulatory cycle phases (Figure 6C). Jacobian analysis of vector fields can help us study cellular state-dependent gene interactions, with positive and negative values of the Jacobian matrix reflecting activation and inhibition regulation, respectively (23). Across all cells, Jacobian analyses demonstrate the activation of *CCNE2* by *CENPE*, repression of *CENPE* by *CCNE2*, self-activation of *CENPE*, and self-activation of *CCNE2* (Figure 6D). These results collectively suggest that the interlinked positive and negative feedback loops between *CCNE2*, which regulates the G1-S checkpoint, and *CENPE*, which regulates the G2-M checkpoint, underlies the dynamics of cell cycle oscillations, revealing the critical role of two transcription factors in maintaining cell cycle progression (Figure 6F). Although only one negative feedback loop is required for biological oscillation, we found that *CCNE2* and *CENPE* have additional self-activating positive feedback loops that promote modularity of cell cycle frequency while essentially unchanged amplitude (Figures 2J and S6). Previous computational studies have shown that it is often difficult to adjust the frequency of a negative feedback oscillator without affecting the amplitude, whereas with positive plus negative feedback, a widely adjustable frequency and a near-constant amplitude can be achieved (69). This also reveals that keeping the cell cycle more robust and easier to evolve requires a lot of positive and negative feedback coupling. To further quantitatively understand the feedback regulation mechanism between regulators, we first plotted distributions of the four Jacobian elements versus expression of each gene (Figure S11A-S11D). Then, we estimated kinetic parameters with derivatives of a simplistic inhibitory or activation hill equation by fitting the Jacobian vs. expression curve in the response heatmap (Figure 6E). Finally, we theoretically and numerically simulate the velocity dynamics curve of the hill equation as a function of gene expression (Figure 6G). The GRN inferred above not only indicates the presence or absence of the interaction between genes but does provide quantitative information on positive and negative regulation relationships.

Similarly, based on the REP1 cell cycle regulation network inferred from scEU-seq data, we further revealed the feedback mechanism between *CCNE1* (a G1-S phase regulator) and *KIF23* (a G2-M phase regulator), and quantified the intensity of feedback between each other (Figures S11E-S11H and S12). By inferring the feedback mechanism between different cell cycle phase regulators, we confirmed that the cell cycle regulatory network is composed of motifs coupled with many negative feedback loops, which together constitute a topology that maintains periodic oscillation. These analyses not only infer the regulatory network of cell cycle function from single-cell omics data, so as to find out the topological basis of cell function, but also further quantify the interaction strength between the regulatory factors, which can provide newly quantitative and accurate guidance for the on-demand design of functional elements in synthetic biology (70).

## Discussion

Spectacular advances in experimentation during the last decade, for example, single-cell dynamics in microscopy and in high throughput data acquisition, have yielded an unprecedented wealth of data on cell dynamics, genetic regulation, and organismal development. These experimental data have motivated the development and refinement of concepts and tools to dissect the physical mechanisms underlying biological processes. An appropriate analytical framework is needed to use these data to study the underlying mechanisms through the non-equilibrium dynamics and thermodynamic behavior of cells. Our study utilizes high-throughput sequencing experimental data to verify the landscape and flux theory and provides its associated quantifications. The cell cycle is a complex process that is tightly regulated to ensure proper cell growth, differentiation, and proliferation. Dysregulation of the cell cycle can lead to various diseases, including cancer. Therefore, understanding the fundamental mechanisms that govern the cell cycle is critical for advancing our knowledge of cellular physiology and developing therapeutic strategies for treating diseases. In this study, we utilized an integrative approach combining single-cell transcriptomics with nonequilibrium statistical physics to better understand the complex regulatory networks and analyze the molecular events involved in the cell cycle at the single-cell level to fill this unmet gap. This process includes processing the upstream dimension reduction clustering of single-cell transcriptome data of the cell cycle, estimation of RNA velocity, reconstruction of cell cycle vector fields, and quantification of non-equilibrium dynamic and thermodynamic landscapes-flux.

Conventionally, researchers have focused on examining the dynamics of trajectories or analyzing the local stabilities of the nonlinear dynamical systems. However, these approaches often offer only a local view of the system’s stability and behavior. To obtain a more comprehensive understanding of the overall stability and behavior, people have turned to the concept of landscape, which can describe the global stability of the system. However, it has become apparent that relying solely on the landscape for the dynamics is true only for equilibrium systems with detailed balance (no net input to or output from the system) is insufficient for capturing the entire dynamics for the nonequilibrium systems (7–9, 17, 43, 71, 72). Our findings in this study of the cell cycle illustrate this point well, as its complex behavior requires the presence of an additional rotational flux. While the landscape contributes to attracting the system into the close oscillation ring valley, it is the flux that drives the coherent oscillation along the cycle path. By combining both concepts, one can gain deeper insights into the mechanisms behind the stability and behavior of the complex systems.

One of the key findings of our research is the significant impact of the biological noise on the underlying mechanism of the cell cycle via nonequilibrium dynamics and thermodynamics. In the small noise range, when the cell cycle oscillation is not disrupted, the noise increases the cell cycle amplitude while conversely decreases the cell cycle period, accompanied by the decrease in the curl flux and EPR. Larger flux means more energy input and larger cell cycle driving force and therefore lead to less time to complete the cell cycle oscillation (the period is smaller). Increased the noise effectively makes potential landscape basins shallower, thereby broadening the distribution of biological variables. After the noise increases to a certain extent, the potential energy landscape shows completely different basins, the cell cycle is broken, and the cells arrest in different cycle stages (attractor-basins). At this time, as the noise increases, although the curl flux increases, it appears that the dominant role of the noise masks the energy supply which is not enough to maintain the coherent oscillation. Our study also demonstrated that the key genetic perturbations can significantly alter the underlying mechanism of the cell cycle via nonequilibrium dynamics and thermodynamics. Specifically, perturbations of cell cycle gene related to cell cycle initiation, DNA replication or mitosis all lead to landscape-flux changes that are consistent with biological effects. These highlight the importance of genetic perturbations in the cell cycle and can be used to predict the pathological effects of certain genetic perturbations, providing a framework for understanding the molecular basis of genetic diseases and diseases that affect the cell cycle. By identifying the bottlenecks and rate-limiting steps in the cell cycle, researchers may be able to develop targeted therapies that can disrupt the cell cycle and selectively kill cancer cells (73). Therefore, future research could build on these findings to develop more accurate models of the cell cycle and other biological processes, and to explore the potential impact of genetic and environmental perturbations on these systems.

We also studied the underlying mechanism via non-equilibrium dynamics and thermodynamics of the cell cycle initiation and termination by counting MFPTs and LAPs between different cell cycle phases. In general, MFPT is defined as the average transition time of a system switching from one state to another, which can be measured experimentally accurately (74). The positive correlation between barrier height and MFPT more intuitively shows the reasons for the difficulties of cell initiation and termination from the 3D landscape, and it is worth noting that our previous works can experimentally quantify the landscape and barrier heigh (74–76). The analysis of EPR and MFPT reveal the existence of basal energy supply in the cell cycle, as well as the initial energy dissipation for cell cycle initiation (44), which approximates the product of EPR and MFPT from G0 phase to G1-S phase. We also found that the cell cycle is a highly energy-dissipative process, and the maintenance of the cell cycle requires thermodynamic energy-dissipation and dynamic flux to enhance the coherence of the oscillation phase against stochastic fluctuations. From a thermodynamic point of view, living systems are open systems in a non-equilibrium state, requiring material input and energy consumption to maintain their steady state and order (77, 78). The Thermodynamic Uncertainty Relationship is a result in non-equilibrium statistical mechanics that relates the dissipation of energy or entropy production to the precision of a thermodynamic observable. In systems far from equilibrium, there is a trade-off between the dissipation of energy (or entropy production) and the fluctuations in the observed quantities. The TUR establishes a lower bound on the fluctuations of a thermodynamic observable based on the rate of entropy production or energy dissipation in the system. The cell cycle is naturally a living system, G0 phase is a non-dividing state of the cell cycle, in which the cell is metabolically active but not actively dividing. To switch from G0 phase to the G1/S phase, the cell requires a certain amount of energy to initiate the complex set of biochemical reactions that drive cell division. Once the cell has entered the G1/S phase, it must maintain a complex system of oscillations and feedback loops in order to progress through the rest of the cell cycle. This requires additional energy input to maintain the oscillations of key cell cycle regulators such as cyclins and cyclin-dependent kinases (CDKs). Currently, certain experiments can quantify the energy flow by measuring the rate of oxygen consumption or heat production or the fluxes of metabolites including the production and comsumption of ATP (79).

In systems biology, the creation of core network regulation to analyze biological processes is an important and popular topic. In general, there are two ways to build a biological regulatory network: one is to create a small network based on the existing knowledge and database, and use the simulator to improve the network, but it is inefficient and cannot be used to build a new network model (80, 81). The second is to use bioinformatics technology, especially high-throughput sequencing data, to infer genes and gene correlations, but to ignore the actual biological modulation relationship (82). To balance the advantages and disadvantages of these two approaches, we infer the transcriptional regulatory networks based on differentially expressed genes of each cycle phase from single-cell transcriptomics data. The regulatory direction between genes (activation/inhibition) can be inferred by Jacobian analyses of vector fields and the hill function of the core network can be simulated by fitting Jacobian in the gene expression space and response heatmap. We reveal that cell cycle regulation involve multiple positive and negative feedback loops between these regulators, which help to adjust the frequency of the oscillation without significantly affecting the amplitude, enabling widely adjustable frequencies and near-constant amplitudes and with maintaining balance between the activation and inhibition of the gene expressions, permitting for proper progression through different cell cycle phases. In this regard, further research is required to unravel such complex regulatory networks comprehensively. Our results identify that some dynamic biological processes can be accurately mapped by constructing regulatory network models based on the high-throughput sequencing data.

In summary, we have built a general framework for the quantification of landscape-flux and analysis of the underlying mechanism of the cell cycle via non-equilibrium dynamics and thermodynamics based on single-cell transcriptomic data that can be applied to numerous biological systems. The mathematical model behind RNA velocity quantity describes only the biological transformation from DNA to RNA, which does not fully capture the intricacies of post-translational regulation from RNA to protein and protein phosphorylation. Corresponding to landscape-flux, the flux driving force may become smaller when protein phosphorylation is lacking. It is worthwhile noticing although some details are missing, the main topology of the Mexican landscape and corresponding flux are similar to the protein level studies (9). This implies that the global nature of the cell cycle network may have already been captured at the transcription level. Of course, more information will need to be complemented at the protein level to study the detailed physical mechanisms. For further research in the future, we need to combine proteomics to evaluate protein velocity and even combine epigenomics with chromatin velocity to understand the regulation mechanism of the cell cycle more comprehensively. More broadly, when coupled with remarkable advances in single-cell approaches, including multi-velocity (83), construction of vector fields (35), as well as spatial multi-omics (84), we will enable a deeper, quantitative understanding of the spatiotemporally underlying mechanism of the cell cycle and other biological systems via non-equilibrium dynamics and thermodynamics to help prevent various diseases, including cancer and neurodegeneration, and open up new possibilities for precision medicine.

## Materials and methods

### Single-cell sequencing data processing and analysis

1. For the scRNA-seq data for ~1k U2OS-FUCCI data, we followed the snakemake pipeline at https://github.com/CellProfiling/FucciSingleCellSeqPipeline to perform scRNA-Seq data preparation and general analysis including filter, dimensionality reduction, clustering analysis and RNA velocity analysis (18).
2. For the scEU-seq data for ~3k RPE1-FUCCI cells, we used *dyn.sample_data.scEU_seq_rpe1* to acquire the processed data by using Dynamo (23), which includes cell cycle clustering and RNA velocity.
3. We reconstructed the vector field based RNA velocity with *dyn.vf.VectorField* by using Dynamo (23). Then, we calculated the divergence, curl, acceleration and curvature to perform the differential geometry analysis. We also calculated the jacobian to perform genetic perturbation and inference gene regulatory interaction.
4. We learned an analytical function of vector field from sparse single cell samples on the entire space robustly by *vector_field_function.* Then, we calculated stochastic dynamics by solving the Langevin equation based analytical function and quantified the non-equilibrium thermodynamics and dynamics of the cell cycle.

### RNA velocity

RNA velocity calculations using Velocyto (21), Scvelo (22) and Dynamo (23). In brief, after the gene selection and normalization, the first- and second-order moments were calculated with *scv.pp.moments* function. The full splicing kinetics were recovered with *scv.tl.recover_dynamics* function and the velocities were obtained with *scv.tl.velocity* function in dynamical mode. The velocities were projected onto diffusion maps and visualized as streamlines with *scv.pl.velocity_embedding_stream* function. The spliced vs. unspliced phase portraits of individual genes were visualized with *scv.pl.velocity*. The pseudotimes of cells were obtain with *scv.tl.recover_latent_time* function.

### Reconstructing vector field from RNA velocity

Dynamo (23) was used to reconstruct function of the continuous and analytical velocity vector field from sparse, noisy single-cell velocity measurements with sparse approximation of vector field consensus (sparseVFC)(87). SparseVFC uses a vector-valued kernel method built on reproducing kernel Hilbert space (RKHS) to learn the vector field, which is expressed analytically as a weighted linear combination of a set of vector-valued kernel basis functions. The learning process relies on sparse approximation to estimate the coefficients (weights) of a selected number of basis functions, each associated with a control point, which is often much smaller than the number of data points. With sparse approximation, the reconstruction scales linearly with the number of data points in both computational time and memory requirements. To account for the noise and outliers of velocity measurements, sparseVFC relies on an EM algorithm to iteratively optimize the set of inliers, as well as the optimized coefficient set for each basis function, further improving the robustness of vector field reconstruction. With the continuous vector field that is learned in either high-dimensional principal component analysis (PCA) space, which can be projected back to the full transcriptomic space, or lower dimensional space (such as 2D uniform manifold approximation and projection [UMAP] space), or directly in the full gene expression space.

### Quantification of potential landscape

The stochastic dynamics of the cell cycle system is described by Langevin equation:

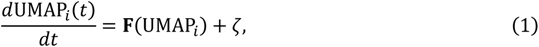

where UMAP*_i_* represents each dimension of UMAP space, which can be projected back to the full transcriptomic space. **F** is the reconstructed vector field function. The noise term (adopts the independent additive white Gaussian noise, 〈ζ(*t*)〉 = 0 and 〈ζ(*t*)((*t′*)〉 = 2*D*δ(*t* − *t′*). *D* is the diffusion coefficient matrix. The noise term is associated with the intensity of cellular fluctuations either from environmental external fluctuations or intrinsic fluctuations. Under large n expansions, the process follows Brownian dynamics. To obtain the potential landscape of cell cycle dynamics, starting from enough random initial conditions, the system will eventually evolve to different cell cycle phase steady state, which may be monostable, multi-stable or limit cycle steady state. All steady-state trajectories in the UMAP space are divided into several small regions, and the steady-state trajectory density in each small region is counted as steady-state probability *P_ss_*. It includes the effects of diffusion through the resulting steady-state probability distribution. U is the generalized potential related to the steady-state probability by U = − ln *P_ss_*.

### Driving force decomposition to the gradient of the potential landscape and curl flux

The probabilistic evolution of diffusion equation, ∂*P*/∂*t* + ∇ · **J**(**x**, *t*) = 0, represents a conservation law of probability (local change is due to net flux in or out). And the probability flux vector **J** of the system in concentration space **x** is defined as 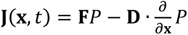. In general, the dynamic driving force **F** can be decomposed into a gradient of a potential and a curl flow flux (7):

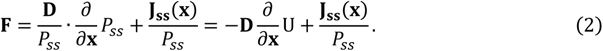

*P_ss_* represents steady state probability distribution and potential landscape U is defined as U = − ln *P_ss_*. At steady state, 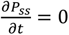, thus ∇ · **J***_ss_* = 0. With detailed balance, there is no net flux and **J***_ss_* = 0, the gradient of potential controls the underlying dynamics as the driving force. For nonequilibrium systems, although ∇ · **J***_ss_* = 0, the net flux **J***_ss_* ≠ 0. Therefore, both the gradient of potential landscape and nonzero steady state probability net flux of probability together form the driving force and determine the dynamics and global properties. The dynamics of a nonequilibrium network can be described as a spiral, along the gradient direction, not as the case of the equilibrium state only following the gradient. It is similar to the electrons moving in both electric and magnetic fields (7).

### EPR (Entropy Production Rate)

In a nonequilibrium open system, there are constant exchanges in energy and information. This results in the dissipation of energy. The dissipation gives a global physical characterization of the nonequilibrium system. In the steady state, the dissipation of energy is closely associated with the entropy production rate. The entropy formula for the system is well known (88),

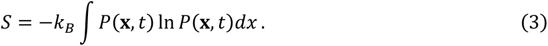

By differentiating the above function, the increase of the entropy at constant temperature *T* is shown as follows:

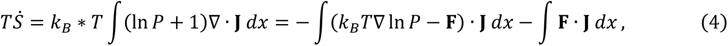

where − ∫(*k_B_T*∇ ln *P* − **F**) · **J** *dx* = EPR is the entropy production rate, and ∫**F** · **J** *dx* = *HDR* is the mean rate of the heat dissipation. In steady state, *Ṡ* = 0, and the entropy production rate EPR is equal to the heat dissipation rate HDR. In this article, we calculated the heat dissipation rate at steady state and also verified that it is the same numerically as entropy production rate at steady state.

### Flux Integration

To quantify the curl probabilistic flux, we define *Flux_Loop_* as the flux integration along the loop trajectory (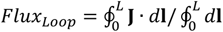) of the cell cycle oscillation divided by the loop length in gene expression space. From **J** = **F***P* − **D** · ∇*P*, we have 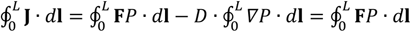. We calculated the flux integration results in 2-dimensional UMAP space projection.

### *In silico* perturbation

For genes in cells, genetic perturbations affect their velocity and thus the changes in the vector field. For small perturbations (Δ**x**) of the gene expression in cells, specific to the influence of perturbation dx_1_, dx_2_, …, dx_n_ of each gene on velocity or vector field, it can be calculated with the exact differential:

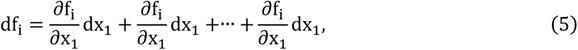

In vectorized form:

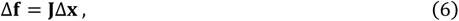

**J** is the Jacobian matrix of vector field and Δ**F** is the velocity and vector field response *in silico* genetic perturbation. In practice, a proportionality constant c is often added to the perturbation Δ**x** to amplify the response Δ**F**.

We first used the *dyn.vf.perturbation* function to perform *in silico* perturbation and visualize the RNA velocity from the perturbation effect vectors. Then, reconstruct the perturbed vector field from the perturbed RNA velocity with *dyn.vf.vectorfield* function. Finally, we perform random dynamics simulation on the vector field of the genetic perturbation to obtain the perturbed landscape, and then quantify the relevant flux and EPR.

### LAPs (Least Action Paths)

Considering a stochastic dynamical system described by Langevin equation:

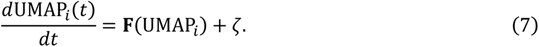

Following the approaches based on the Freidlin-Wentzell theory (89), the most probable transition path from attractor j at time 0 to attractor *k* at time *T*, can be acquired by minimizing the action functional over all possible paths:

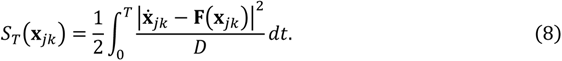

Where *D* is the diffusion coefficient accounting for the stochasticity of gene expression, and for simplicity here we assume it to be a constant. This path **x** is called the least action path (LAP). Specifically, the optimal path between any two cell states is searched by variating the continuous path connecting the source state to the target while minimizing its action and updating the associated transition time. The resultant LAP has the highest transition probability and is associated with a particular transition time. This type of approach has been used in many chemical and biological systems.

### MFPT (Mean First Passage Time)

For rare transitions with *S_T_* ≫ 0, the transition rate (number of transitions per unit time) is proportional to the exponential of the actions. The Freidlin–Wentzell theorem dictates that the LAP with the minimal traversal time (which will be referred to as the optimal path below) contributes the most to this transition rate (89):

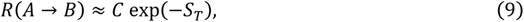

where *A* and *B* are two cell cycle phases, *S_T_* is the action of the optimal path, and *C* a proportional factor. Furthermore, the transition time, or more specifically the mean first passage time (MFPT), is related to the transition rate:

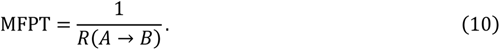

Therefore, the action of the optimal path predicts both the likelihood and transition time for such rare transitions.

### Phase diffusion and coherence time

We simulated 500 trajectories starting with the same initial condition. For the *j*-th trajectory, we obtained its *i*-th peak time *t_i_* from the trajectory **x_J_**(*t*) after smoothing (smooth function in MATLAB was used). The peak positions for two trajectories are shown in Figure S5. For each of the trajectories, we computed the mean of their *i*-th peak time *m_i_* = ∑*_j_ t_ij_*/*N*, and variance 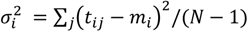, where *N* is the total number of trajectories. The average period *T* is given by *T* = *m_i_*/*i*. Asymptotically, 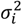 depends linearly on *m_i_*, and the slope of this linear dependence is the peak-time diffusion constant *D*, which has the dimension of time. The phase diffusion constant *D*_∅_ is linearly proportional to *D* (47):

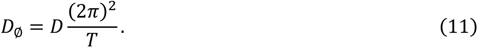

The autocorrelation function (47):

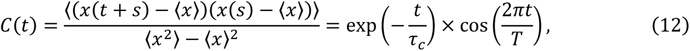

where *s* is a time variable, the period is given by *T* and R*_C_* is the coherence time for the oscillation.

### Power Spectral Density (PSD) from autocorrelation

The distribution of average power of a signal **x**(*t*) in the frequency domain ω is called the power spectral density (PSD) or power density (PD) or power density spectrum. In order to drive the PSD function, consider a power signal as a limiting case of an energy signal, i.e., the signal *Z*(*t*) is zero outside the interval |τ/2|, the signal *Z*(*t*) is given by,

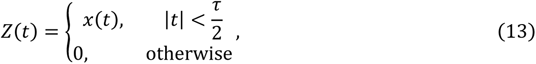

Where, **x**(*t*) is a power signal of same magnitude extending to infinity.

As the signal *Z*(*t*) is finite duration signal of duration *τ* and thus, it is an energy signal having energy E, that is given by,

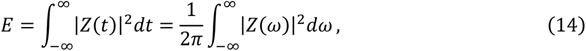

Where 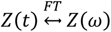. Also,

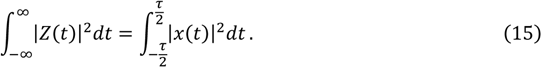

Therefore, we have,

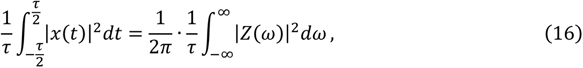

Hence, when τ → ∞, then the LHS of the above equation gives the average power (P) of the signal *x*(*t*), i.e.,

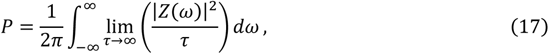

if *τ* → ∞, then 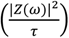 approaches a finite value. Assume this finite value is represented by *S*(ω), i.e.,

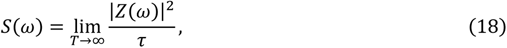

this expression is called the power spectral density (PSD) of the signal *Z*(*t*). Therefore, for the function *x*(*t*), the PSD function is given by,

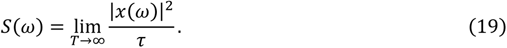

The power spectral density function *S*(ω) and the autocorrelation function *R*(*τ*) of a power signal form a Fourier transform pair, i.e.,

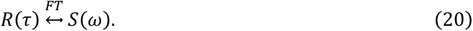

### Phase coherence

The robustness of the oscillation with respect to the diffusion coefficient *D* can be quantified further by the phase coherence ξ, which measures the degree of periodicity of the time evolution of a given variable (49). The phase coherence ξ quantitatively measures the degree of persistence of the oscillatory phase and is defined as follows. First, the vector *N*(*t*) = *n*_1_(*t*)*e*_1_ + *n*_2_(*t*)*e*_2_ is shown in Figure S10I. The unit vectors are *e*_1_ = (1,0) and *e*_2_ = (0, 1), and *n*_1_(*t*) and *n*_2_(*t*) are the concentration of the two kinds of protein molecules at time *t*. Then, *ϕ*(*t*) is the phase angle between *N*(*t*) and *N*(*t* + *τ*), where *τ* should be smaller than the deterministic period and larger than the fast fluctuations. *ϕ*(*t*) > 0 to represent that the oscillation goes on the positive orientation (counterclockwise). The formula of ξ is shown as

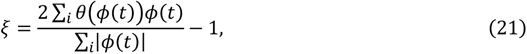

where *θ*(*ϕ*) = 1 when *ϕ*(*t*) > 0, and 0(*ϕ*) = 0 when *ϕ*(*t*) ≤ 0, and sums are taken over every time steps for the simulant trajectory. ξ ≈ 0 means the system moves stochastically and has no coherence. The oscillation is most coherent as ξ is close to 1.

### Gene regulatory network inference

To construct genes interaction network of the cell cycle, we first calculated and obtained the all differentially expressed genes in different cell cycle phase (Tables S1 and S2). After that the interaction between the differentially expressed genes were calculated by Jacobian based on the reconstructed vector field function. Positive values from Jacobian represent activation and negative values represent inhibition (23). The visualization of all networks was performed using Cytoscape (86). The nodes and edges each represent the gene and interactions in a network.

### Estimating kinetic parameters by fitting the Jacobian vs. expression curve

The Hill function is a mathematical model commonly used to describe various regulatory interactions in genetic and biochemical networks. It is used to represent the relationship between the concentration of a regulator (e.g., transcription factor) and the rate of transcriptional activation or repression. Its versatility lies in its ability to capture a wide range of phenomena and dynamics commonly observed in biological systems. The Hill function for quantifying the gene regulations works reasonably well when regulating binding/unbinding is relatively fast compared to protein synthesis and degradation. Derivation from the Hill function is expected when the regulatory binding/unbinding is relatively slow or comparable compared to protein synthesis and degradation (90, 91). Because the reconstructed vector field is expressed as a set of implicit basis functions, not explicitly as Hill functions, in the current framework we are not able to directly obtain kinetic parameters such as the Hill coefficient. Nevertheless, the reconstructed vector field encodes such information, and additional computations are applied to extract that information (23). We demonstrate this possibility on simplistic network motifs such as CCNE2-CENP2 and CCNE2-KIF23, by fitting the derivatives of inhibitory or activation Hill equations to the corresponding Jacobian elements. Further efforts will be needed to make such efforts generally applicable to systems with more sophisticated mechanisms.

Formally, we assume that the activation effect of gene x on the target gene y takes the form of an activating Hill function:

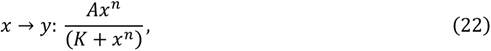

and that the inhibition effect assumes the form of an inhibitory Hill function:

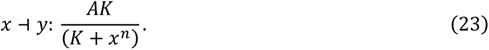

For the activation-inhibition and self-activation of two gene x and y, the ODEs can write as:

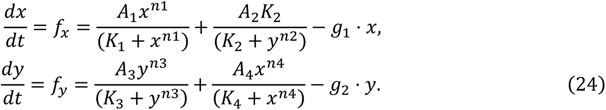

The Jacobian of this system is a 2-by-2 matrix:

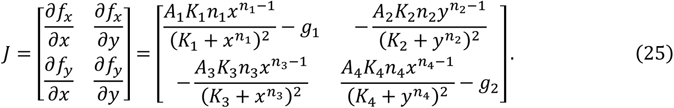

In Figures 6E and S12E, the means and standard deviations of the Jacobian vs. expression profiles were calculated and fitted with the above derivatives using the Dynamo: *dyn.pl.response*, which can fit and obtain the parameters of hill function (Table S3).

### Quantification and statistical analysis

The calculation of the amplitude and period of cell cycle oscillations was averaged multiple times using MATLAB. All correlation analyses and confidence intervals are calculated in Origin. The fitting evaluation of phase diffusion was performed by MATLAB.

## Acknowledgements

Ligang Zhu thanks the supports from the National Natural Science Foundation of China Grant No. 12234019, 21721003. We would also like to thank Kun Zhang, Li Xu, Wenbo Li, Han Yan, Cong Yu, Linqi Wang and Songlin Yang for valuable discussions during the preparation of this manuscript.

## Author contributions

L.Z. designed the research, developed the methods, carried out the stimulations, performed the analysis, and wrote the manuscript. J.W. initiated, designed and supervised the research, developed the methods, analyzed the data, and wrote the manuscript.

## Supplemental information

Supplementary Table S1-S3 and Supplementary Figure S1–S12.

## Data and code availability

The scRNA-seq raw data of U2OS-FUCCI cell cycle from the paper by Mahdessian et al (18), which are available at GEO with accession GSE14677318. The scEU-seq data of RPE1-FUCCI cell cycle can be extracted using dynamo’s CLI: *dyn.sample_data.scEU_seq_rpe1* or from the paper by Battich et al (19), which are available at GEO with accession number GSE12836519. The processed data are available from Github (https://github.com/Zhu-1998/cellcycle). The source codes used in this study is publicly available from Github (https://github.com/Zhu-1998/cellcycle). Requests for additional program codes and data generate and/or analyzed during the current study should be fulfilled on reasonable request by the Lead Contact.

## Supplementary Information

### Supplementary Tables

**Table S1.** Differential expression genes for U2OS cell cycle regulation, related to Figure 6.

| test_group | gene | versus_group | exp_frac | log2_fc | pval | qval |
| --- | --- | --- | --- | --- | --- | --- |
| **G1-S** | *CCNE2* | S | 0.798898 | 1.023973 | 2.64E-16 | 1.33E-15 |
|  | *MSH6* | G2-M | 0.997245 | 1.013831 | 1.13E-63 | 9.91E-62 |
|  | *MCM10* | G2-M | 0.942149 | 1.043559 | 6.55E-37 | 5.05E-36 |
|  | *MCM3* | G2-M | 0.997245 | 1.0391 | 1.85E-71 | 4.84E-69 |
|  | *CDC6* | G2-M | 0.964187 | 1.027817 | 4.11E-61 | 1.54E-59 |
|  | *SLBP* | G2-M | 0.983471 | 1.212273 | 5.23E-60 | 1.25E-58 |
|  | *UNG* | G2-M | 0.972452 | 1.619185 | 2.23E-62 | 9.73E-61 |
|  | *HELLS* | G2-M | 0.936639 | 1.178466 | 1.32E-24 | 8.06E-24 |
|  | *RAD51* | G2-M | 0.895317 | 1.311513 | 9.65E-34 | 7.23E-33 |
| **S** | *MCM6* | G2-M | 0.974576 | 1.21473 | 5.60E-30 | 5.31E-29 |
|  | *SYBU* | G2-M | 0.241525 | 1.409025 | 8.43E-03 | 1.11E-02 |
|  | *RASGEF1B* | G2-M | 0.165254 | 1.180672 | 4.32E-02 | 4.97E-02 |
| **G2-M** | *PSRC1* | G1-S | 0.95 | 1.155866 | 1.27E-51 | 2.04E-50 |
|  | *CCNF* | G1-S | 0.954167 | 1.222709 | 2.62E-47 | 3.31E-46 |
|  | *KIF18A* | G1-S | 0.920833 | 1.06701 | 4.85E-35 | 3.80E-34 |
|  | *FAM83D* | G1-S | 0.941667 | 1.360211 | 6.35E-49 | 8.77E-48 |
|  | *CENPE* | G1-S | 0.920833 | 1.115475 | 4.32E-33 | 2.91E-32 |
|  | *NEK2* | G1-S | 0.929167 | 1.095565 | 1.63E-35 | 1.31E-34 |
|  | *CENPA* | G1-S | 0.9 | 1.013723 | 5.63E-33 | 3.71E-32 |
|  | *GAS2L3* | G1-S | 0.904167 | 1.190276 | 3.53E-24 | 1.50E-23 |
|  | *BORA* | G1-S | 0.904167 | 1.480186 | 4.58E-34 | 3.24E-33 |
|  | *CDC25C* | G1-S | 0.883333 | 1.064069 | 7.03E-22 | 2.68E-21 |
|  | *KIF20A* | S | 0.9125 | 1.027906 | 8.60E-33 | 5.42E-32 |
|  | *FAM72D* | S | 0.516667 | 1.097402 | 1.03E-06 | 2.57E-06 |
| **M** | *PIF1* | G1-S | 0.792593 | 1.813076 | 5.90E-10 | 6.23E-10 |
|  | *H2BC6* | G2-M | 0.37037 | 1.017269 | 2.59E-02 | 2.68E-02 |
| **M-G1** | *PDE4B* | G2-M | 0.27439 | 1.303967 | 7.50E-03 | 7.77E-03 |

**Table S2.**
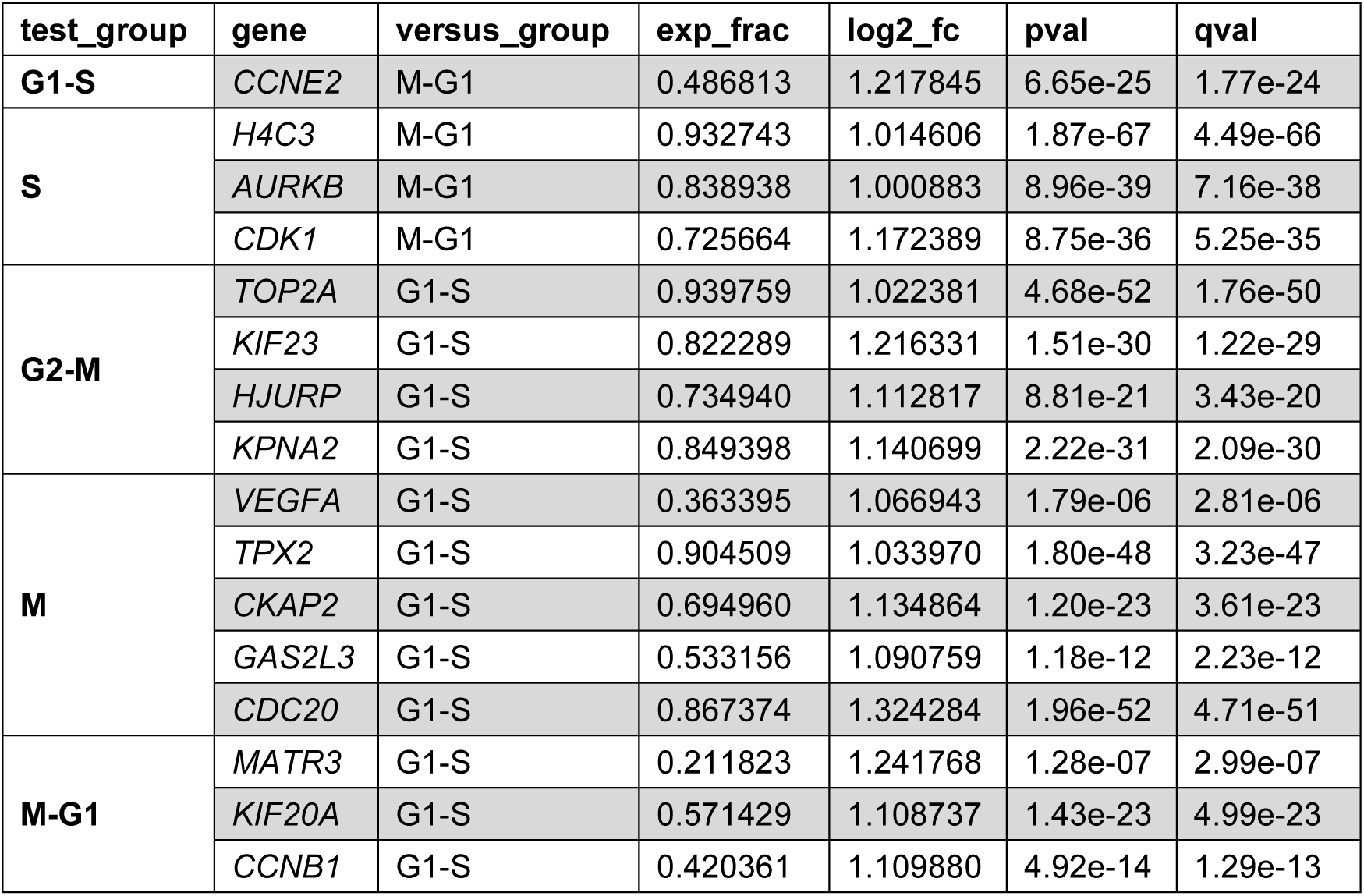
Differential expression genes for RPE1 cell cycle regulation, related to Figure S12.

| test_group | gene | versus_group | exp_frac | log2_fc | pval | qval |
| --- | --- | --- | --- | --- | --- | --- |
| **G1-S** | *CCNE2* | M-G1 | 0.486813 | 1.217845 | 6.65e-25 | 1.77e-24 |
| **S** | *H4C3* | M-G1 | 0.932743 | 1.014606 | 1.87e-67 | 4.49e-66 |
|  | *AURKB* | M-G1 | 0.838938 | 1.000883 | 8.96e-39 | 7.16e-38 |
|  | *CDK1* | M-G1 | 0.725664 | 1.172389 | 8.75e-36 | 5.25e-35 |
| **G2-M** | *TOP2A* | G1-S | 0.939759 | 1.022381 | 4.68e-52 | 1.76e-50 |
|  | *KIF23* | G1-S | 0.822289 | 1.216331 | 1.51e-30 | 1.22e-29 |
|  | *HJURP* | G1-S | 0.734940 | 1.112817 | 8.81e-21 | 3.43e-20 |
|  | *KPNA2* | G1-S | 0.849398 | 1.140699 | 2.22e-31 | 2.09e-30 |
| **M** | *VEGFA* | G1-S | 0.363395 | 1.066943 | 1.79e-06 | 2.81e-06 |
|  | *TPX2* | G1-S | 0.904509 | 1.033970 | 1.80e-48 | 3.23e-47 |
|  | *CKAP2* | G1-S | 0.694960 | 1.134864 | 1.20e-23 | 3.61e-23 |
|  | *GAS2L3* | G1-S | 0.533156 | 1.090759 | 1.18e-12 | 2.23e-12 |
|  | *CDC20* | G1-S | 0.867374 | 1.324284 | 1.96e-52 | 4.71e-51 |
| **M-G1** | *MATR3* | G1-S | 0.211823 | 1.241768 | 1.28e-07 | 2.99e-07 |
|  | *KIF20A* | G1-S | 0.571429 | 1.108737 | 1.43e-23 | 4.99e-23 |
|  | *CCNB1* | G1-S | 0.420361 | 1.109880 | 4.92e-14 | 1.29e-13 |

**Table S3.** The hill function and parameters of gene interaction related to Figures 6 and S12.

| ODEs | Interaction term | Parameters |
| --- | --- | --- |
| The interaction between CCNE2 and CENPE |  |  |
| $\frac{d[CCNE2]}{dt} = f_{CCNE2} = V_1 + V_2 - V_3$ | Self-activation | $A_1 = 3.7\text{e-}3$ |
| | $V_1 = \frac{A_1[CCNE2]^{n_1}}{(K_1 + [CCNE2]^{n_1})}$ | $K_1 = 16.2$<br>$n_1 = 2.7$ |
| | CENPE activates CCNE2 | $A_3 = 9.3\text{e-}3$ |
| | $V_2 = \frac{A_3[CENPE]^{n_3}}{(K_3 + [CENPE]^{n_3})}$ | $K_3 = 83.9$<br>$n_3 = 4.1$ |
| | Degradation<br>$V_3 = g_1 \cdot [CCNE2]$ | $g_1 = 3.4\text{e-}15$ |
| $\frac{d[CENPE]}{dt} = f_{CENPE} = V_4 + V_5 - V_6$ | Self-activation | $A_2 = 1.3\text{e-}2$ |
| | $V_4 = \frac{A_2[CENPE]^{n_2}}{(K_2 + [CENPE]^{n_2})}$ | $K_2 = 37.9$<br>$n_2 = 3.1$ |
| | CCNE2 inhibits CENPE | $A_4 = 9.8\text{e-}4$ |
| | $V_5 = \frac{A_4 K_4}{(K_4 + [CCNE2]^{n_4})}$ | $K_4 = 5.1$<br>$n_4 = 2.6$ |
| | Degradation<br>$V_6 = g_2 \cdot [CENPE]$ | $g_2 = 2.0\text{e-}4$ |
| The interaction between CCNE2 and KIF23 |  |  |
| $\frac{d[CCNE2]}{dt} = f_{CCNE2} = V_1 + V_2 - V_3$ | Self-activation | $A_1 = 4.7\text{e-}4$ |
| | $V_1 = \frac{A_1[CCNE2]^{n_1}}{(K_1 + [CCNE2]^{n_1})}$ | $K_1 = 2.3$<br>$n_1 = 1.1$ |
| | KIF23 inhibits CCNE2 | $A_4 = 3.2\text{e-}4$ |
| | $V_2 = \frac{A_4 K_4}{(K_4 + [KIF23]^{n_4})}$ | $K_4 = 0.9$<br>$n_4 = 3.1$ |
| | Degradation<br>$V_3 = g_1 \cdot [CCNE2]$ | $g_1 = 1.3\text{e-}4$ |
| $\frac{d[KIF23]}{dt} = f_{KIF23} = V_4 + V_5 - V_6$ | Self-activation | $A_2 = 1.5\text{e-}4$ |
| | $V_4 = \frac{A_2[KIF23]^{n_2}}{(K_2 + [KIF23]^{n_2})}$ | $K_2 = 1.7$<br>$n_2 = 6.6$ |
| | CCNE2 activates KIF23 | $A_3 = 4.1\text{e-}4$ |
| | $V_5 = \frac{A_3[CCNE2]^{n_3}}{(K_3 + [CCNE2]^{n_3})}$ | $K_3 = 1.2$<br>$n_3 = 2.4$ |
| | Degradation<br>$V_6 = g_2 \cdot [KIF23]$ | $g_2 = 6.7\text{e-}27$ |

### Supplementary Figures

**Figure S1.**
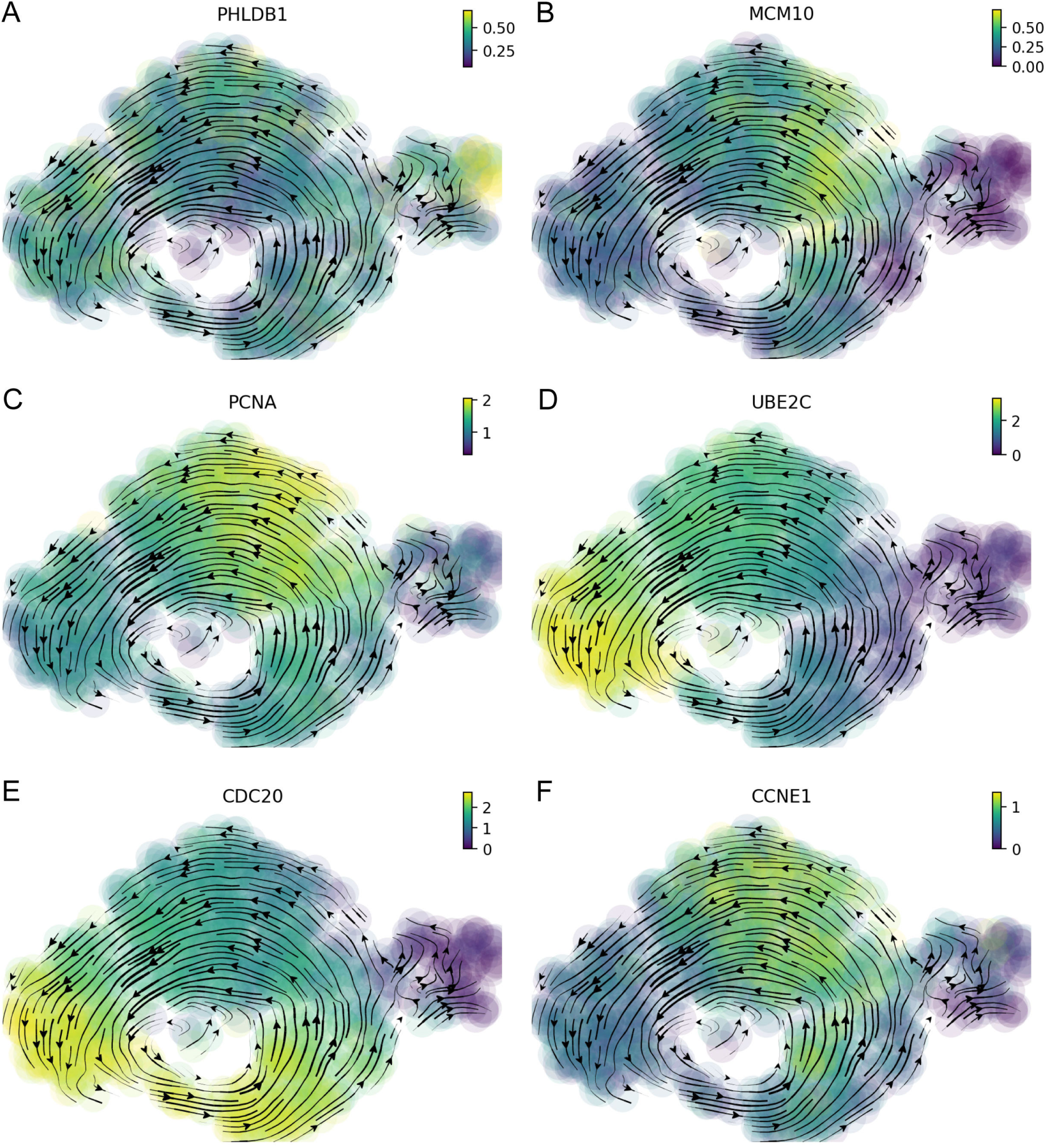
The key gene expression of different cell cycle phases (Related to Figure 1) (A) *PHLDB1* has high expression in the G0 phase. (B) *MCM10* has high expression in the G1-S phase. (C) *PCNA* has high expression in the S phase. (D) *UBE2C* has high expression in the G2-M phase. (E) *CDC20* has high expression in the M phase. (F) *CCNE1* has high expression in the M-G1 phase.

**Figure S2.**
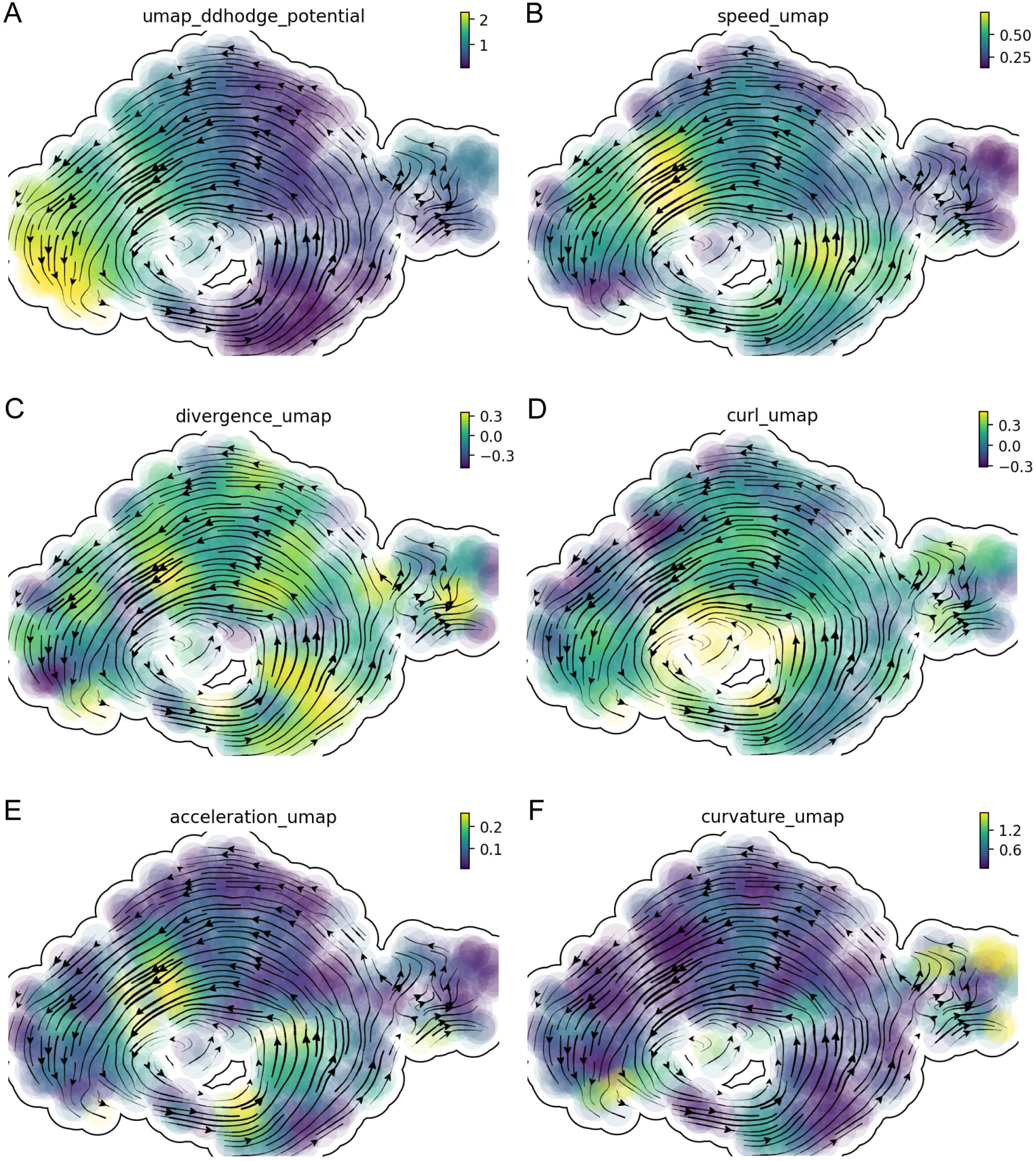
The differential geometric analysis of cell cycle dynamics of U2OS cells (Related to Figure 1) (A) Ddhodge potential of the reconstructed vector field among all cell cycle phases. (B) Same as in (A) but for the speed. (C) Same as in (A) but for the divergence. (D) Same as in (A) but for the curl. (E) Same as in (A) but for the acceleration. (F) Same as in (A) but for the curvature.

**Figure S3.**
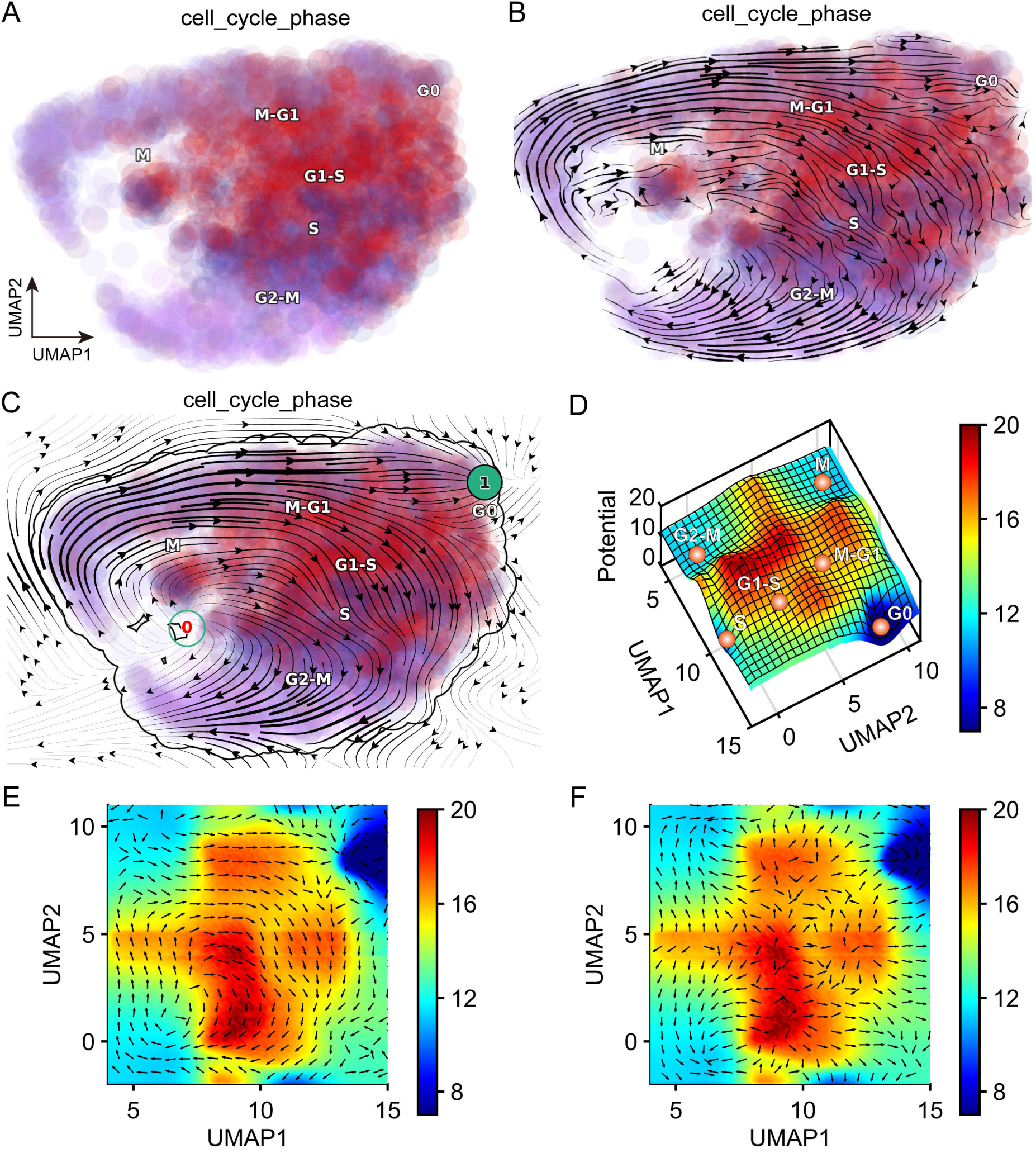
Constructing cell cycle landscape-flux by scEU-seq data of RPE1 cells (Related to Figure 1) (A) Cell cycle phase clusters by UMAP. (B) RNA velocity of cell cycle dynamics. (C) Reconstructed vector field of cell cycle dynamics. (D) Potential landscape of cell cycle dynamics in UMAP. (E) Curl flux (black arrow) of cell cycle dynamics landscape in UMAP. (F) The gradient force (black arrow) of cell cycle dynamics landscape in UMAP.

**Figure S4.**
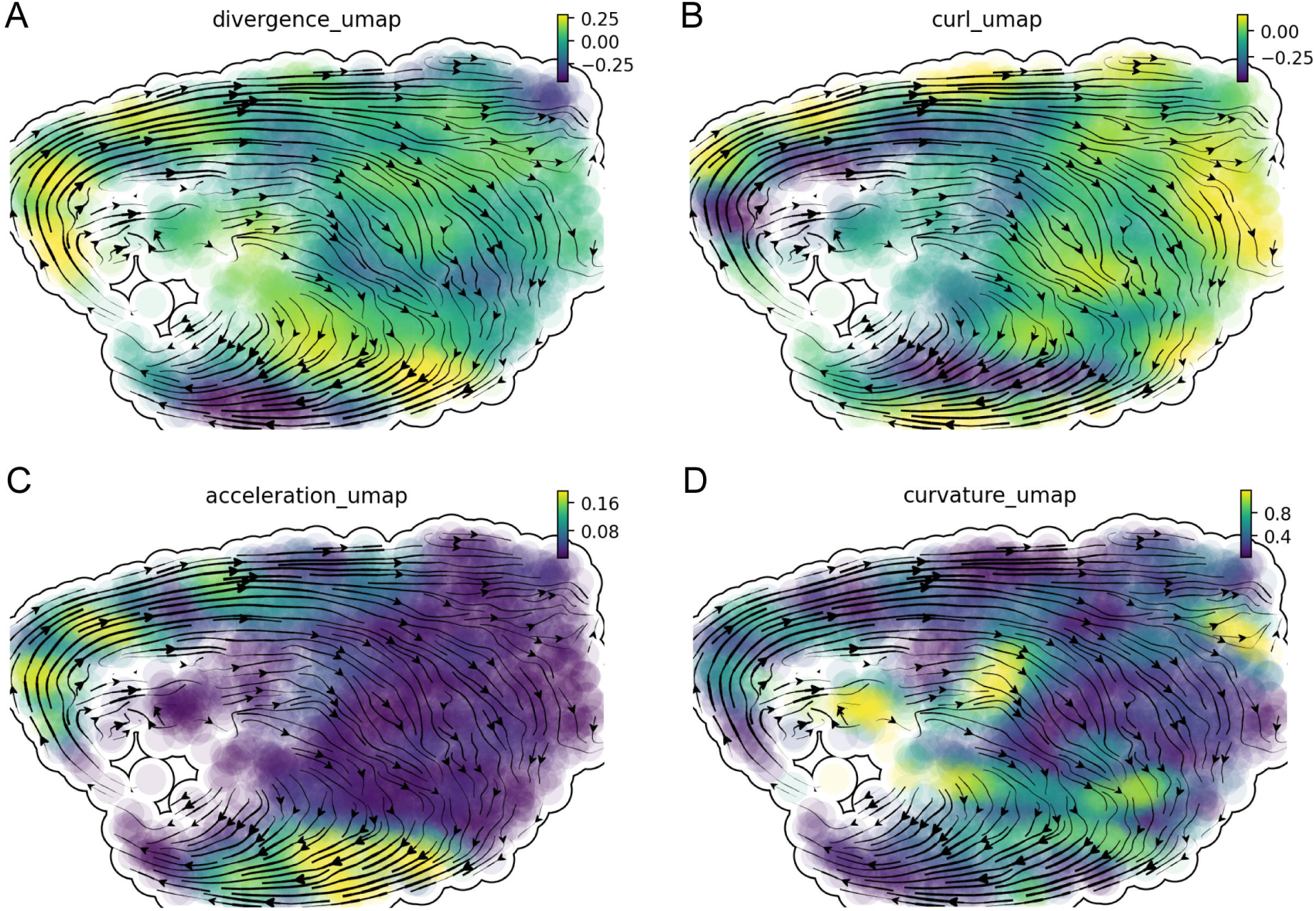
The differential geometric analysis of cell cycle dynamics of RPE1 cell (Related to Figure 1) (A) Divergence of the reconstructed vector field among all cell cycle phases. (B) Same as in (A) but for the curl. (C) Same as in (A) but for the acceleration. (D) Same as in (A) but for the curvature.

**Figure S5.**
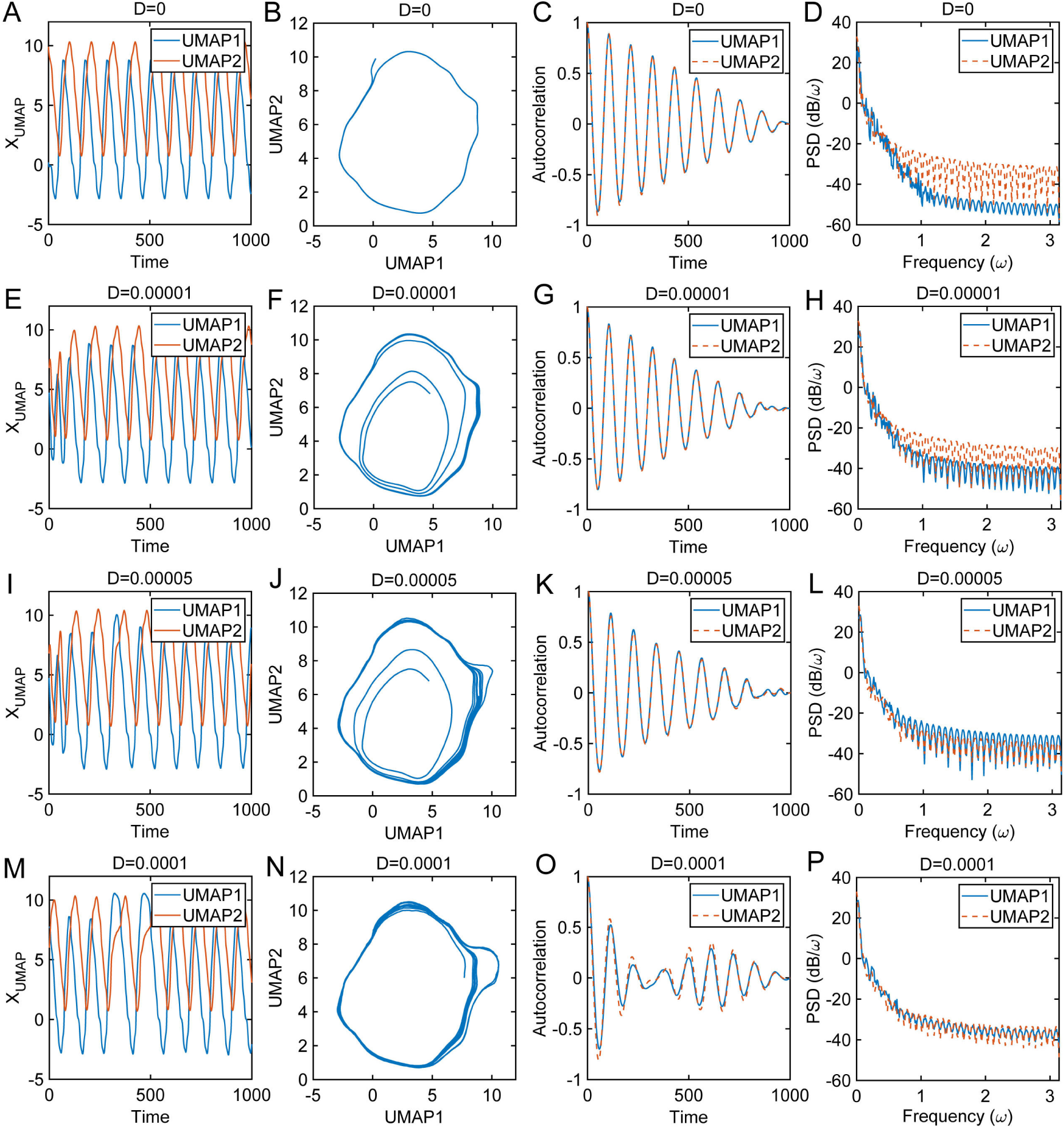
The trajectory, autocorrelation and power spectral density of cell cycle oscillation dynamics with different strength of noise (Related to Figure 2) (A) The trajectories along time of the cell cycle for different diffusion coefficient D=0. (B) The trajectories in phase space corresponding to (A) for different diffusion coefficient D=0. (C) The autocorrelation corresponding to (A) for different diffusion coefficient D=0. (D) The power spectral density corresponding to (A) for different diffusion coefficient D=0. (E) Same as in (A) but for different diffusion coefficient D=0.00001. (F) Same as in (B) but for different diffusion coefficient D=0.00001. (G) Same as in (C) but for different diffusion coefficient D=0.00001. (H) Same as in (D) but for different diffusion coefficient D=0.00001. (I) Same as in (A) but for different diffusion coefficient D=0.00005. (J) Same as in (B) but for different diffusion coefficient D=0.00005. (K) Same as in (C) but for different diffusion coefficient D=0.00005. (L) Same as in (D) but for different diffusion coefficient D=0.00005. (M) Same as in (A) but for different diffusion coefficient D=0.0001. (N) Same as in (B) but for different diffusion coefficient D=0.0001. (O) Same as in (C) but for different diffusion coefficient D=0.0001. (P) Same as in (D) but for different diffusion coefficient D=0.0001.

**Figure S6.**
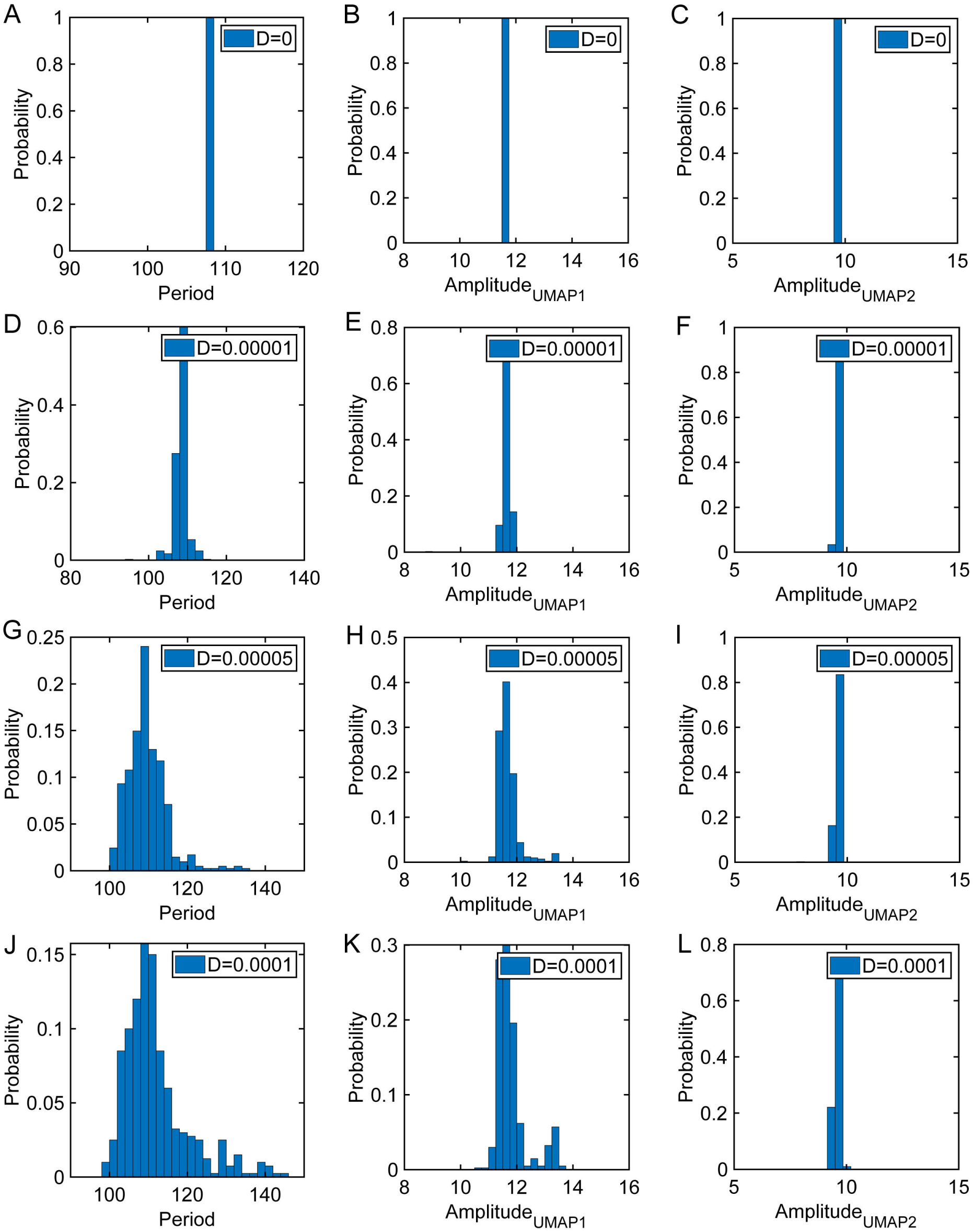
The period and amplitude distribution of cell cycle oscillation dynamics with different strength of noise (Related to Figure 2) (A) The distribution of cell cycle period for different diffusion coefficient D=0. (B) The distribution of amplitude in UMAP1 for different diffusion coefficient D=0. (C) The distribution of amplitude in UMAP2 for different diffusion coefficient D=0. (D) Same as in (A) but for different diffusion coefficient D=0.00001. (E) Same as in (B) but for different diffusion coefficient D=0.00001. (F) Same as in (C) but for different diffusion coefficient D=0.00001. (G) Same as in (A) but for different diffusion coefficient D=0.00005. (H) Same as in (B) but for different diffusion coefficient D=0.00005. (I) Same as in (C) but for different diffusion coefficient D=0.00005. (J) Same as in (A) but for different diffusion coefficient D=0.0001. (K) Same as in (B) but for different diffusion coefficient D=0.0001. (L) Same as in (C) but for different diffusion coefficient D=0.0001.

**Figure S7.**
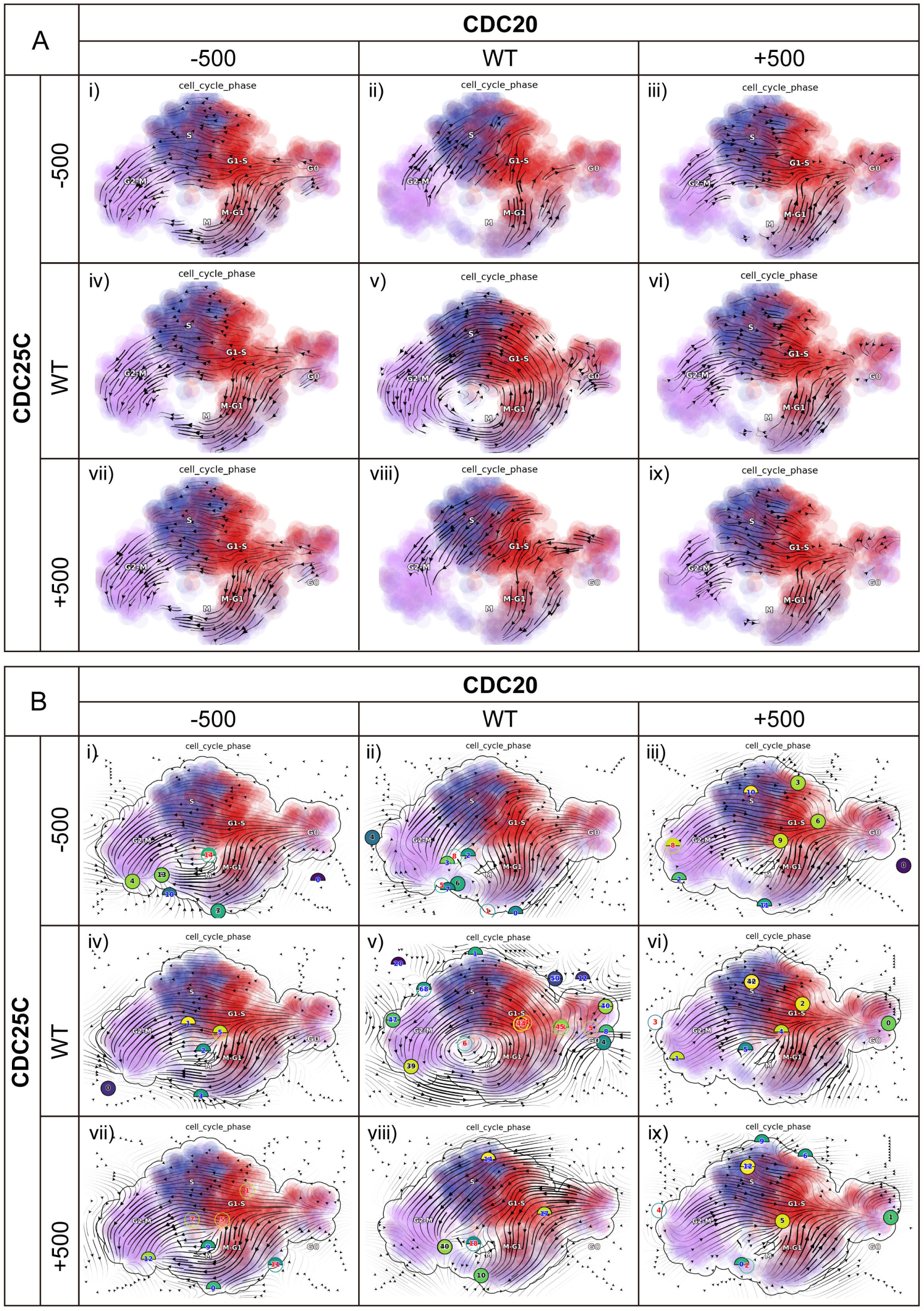
RNA velocity and reconstructed vector field of cell cycle under genetic perturbation (Related to Figure 3) (A) *In silico* perturbation trajectory predictions: (i) Suppression of both *CDC20* and *CDC25C*. (ii) Suppression of *CDC25C* only. (iii) Activation of *CDC20* and suppression of *CDC25C*. (iv) Suppression of *CDC20* only. (v) WT. (vi) Activation of *CDC20* only. (vii) Suppression of *CDC20* and activation of *CDC25C*. (viii) Activation of *CDC25C* only. (ix) Activation of both *CDC20* and *CDC25C*. (B) *In silico* perturbation vector field predictions: (i) Suppression of both *CDC20* and *CDC25C*. (ii) Suppression of *CDC25C* only. (iii) Activation of *CDC20* and suppression of *CDC25C*. (iv) Suppression of *CDC20* only. (v) WT. (vi) Activation of *CDC20* only. (vii) Suppression of *CDC20* and activation of *CDC25C*. (viii) Activation of *CDC25C* only. (ix) Activation of both *CDC20* and *CDC25C*.

**Figure S8.**
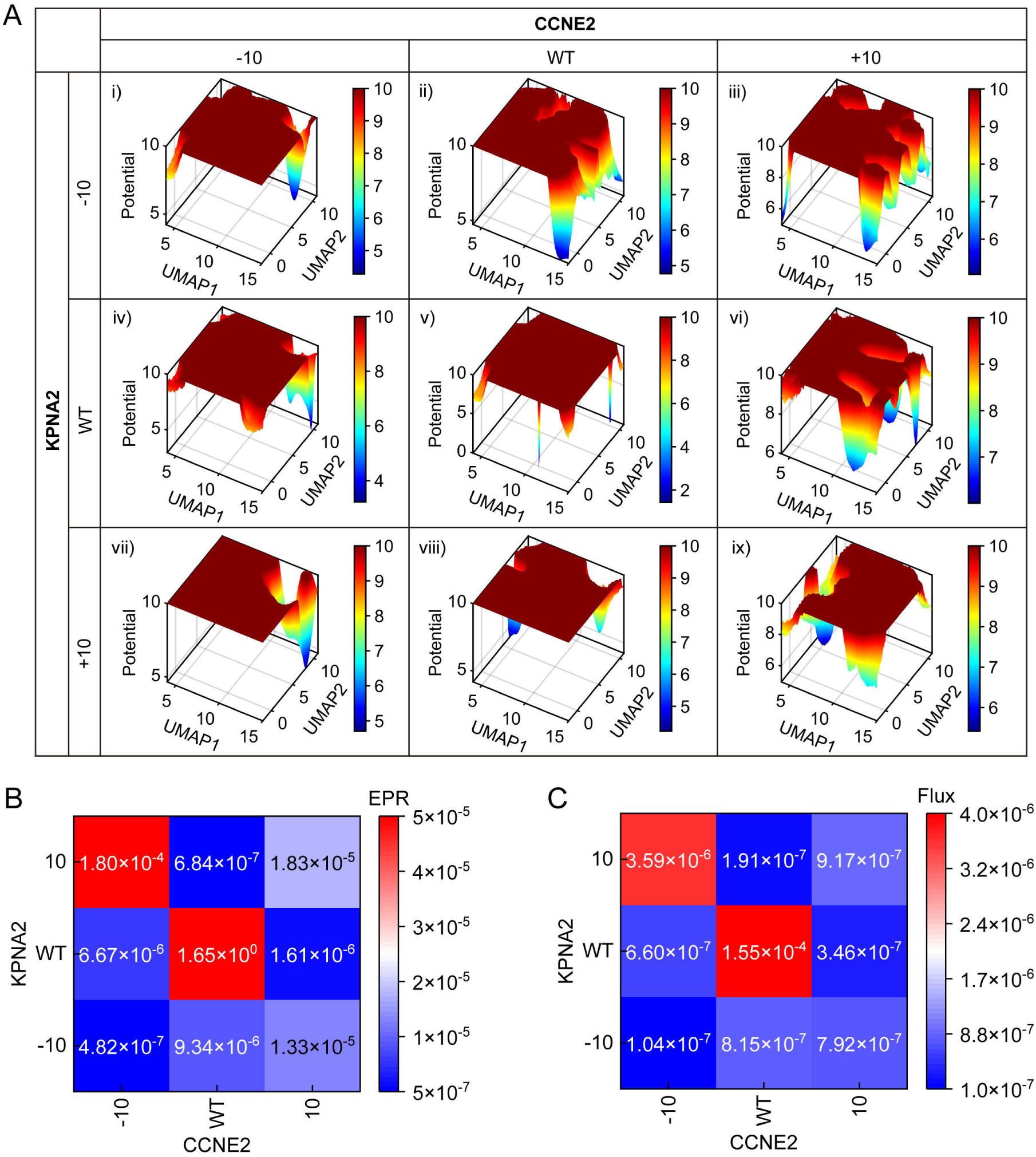
The genetic perturbation alters of RPE1 cell cycle landscape-flux (Related to Figure 3) (A) Cell cycle global dynamics landscape of REP1 cells in UMAP with different genetic perturbation. (i) Suppression of both *CCNE2* and *KPNA2*. (ii) Suppression of *KPNA2* only. (iii) Activation of *CCNE2* and suppression of *KPNA2*. (iv) Suppression of *CCNE2* only. (v) WT. (vi) Activation of *CCNE2* only. (vii) Suppression of *CCNE2* and activation of *KPNA2*. (viii) Activation of *KPNA2* only. (ix) Activation of both *CCNE2* and *KPNA2*. (B) EPR of cell cycle nonequilibrium thermodynamics with different genetic perturbation. (C) The average Flux of cell cycle nonequilibrium dynamics with different genetic perturbation.

**Figure S9.**
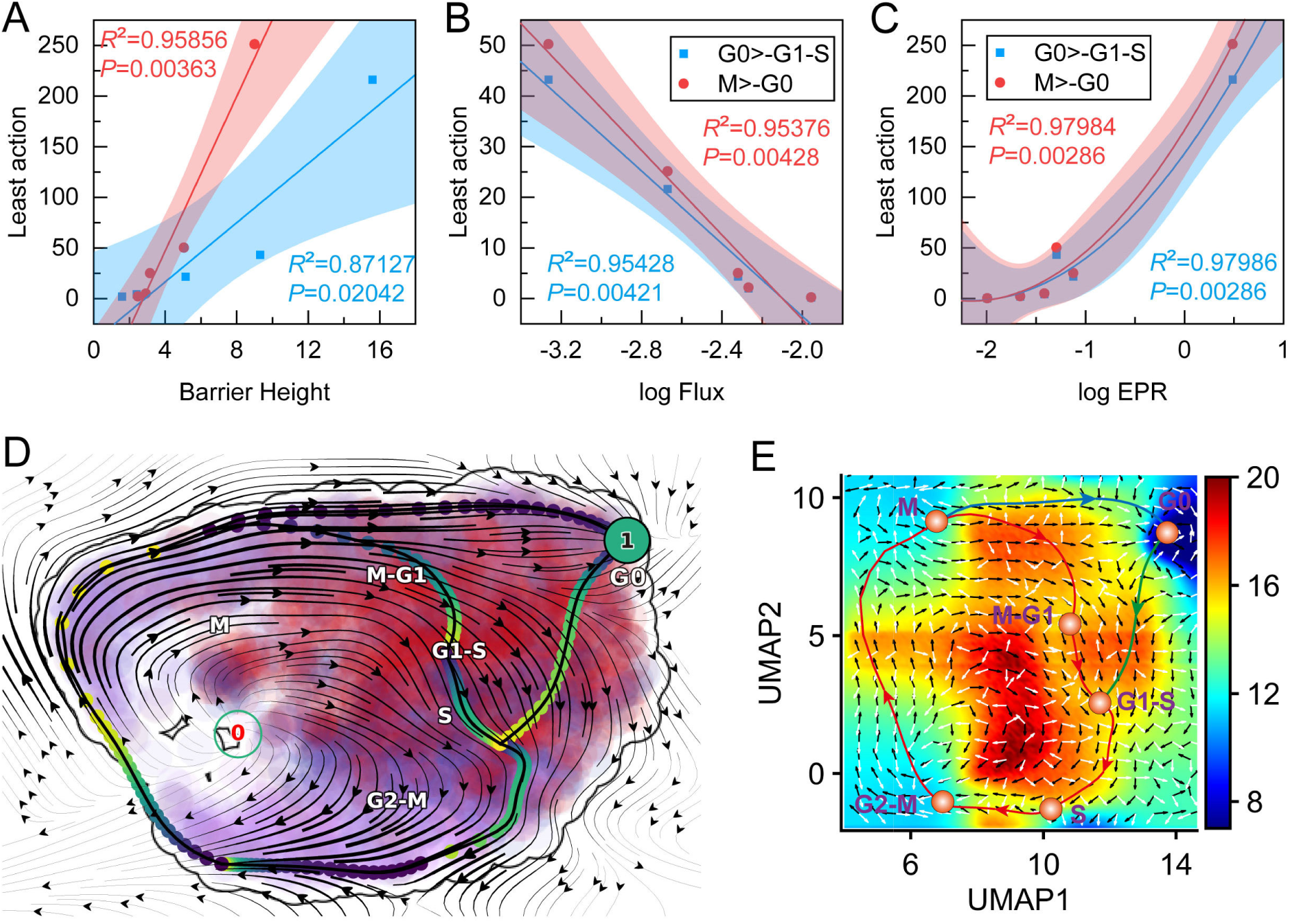
LAPs and MFPT in cell cycle initiation and termination (Related to Figure 4) (A) The correlation between barrier height and least action when diffusion coefficients are changed. The line is a fitting correlation line, and the shaded part is the 95% fitting confidence interval. (B) The correlation between the logarithm of flux and least action when diffusion coefficients are changed. (C) The correlation between the logarithm of EPR and least action when diffusion coefficients are changed. (D) LAPs between different cell cycle phases in the velocity vector field of RPE1 cells. The color of digits in each node reflects the type of fixed point: red, emitting fixed point; black, absorbing fixed point. The color of the numbered nodes corresponds to the confidence of the fixed points. The color of the dots along the paths corresponds to the direction. (E) LAPs between different cell cycle phases in 2D landscape of RPE1 cells. The black arrows present the curl flux and the white arrows present the gradient force in the landscape.

**Figure S10.**
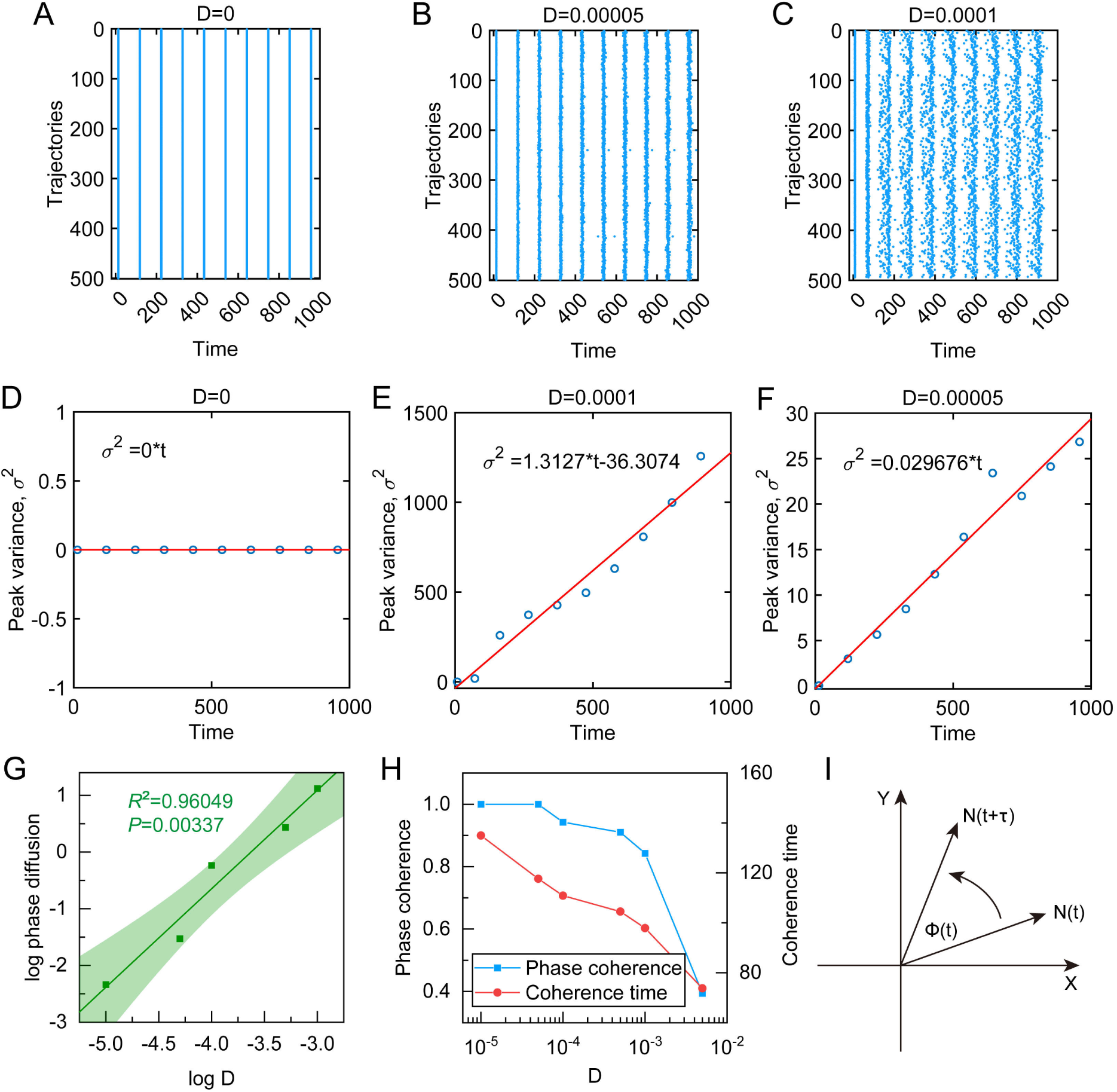
Phase diffusion and phase coherence of cell cycle oscillation with different noise (Related to Figure 5) (A-C) Raster plot of the peak times for 500 different trajectories starting with the same initial condition at the different diffusion coefficient, D=0 (A), D=0.00005 (B) and D=0.0001 (C). (D-F) Peak time variance σ^2^ goes linearly with the average peak time for the different diffusion coefficient, with the linear coefficient defined as the peak-time diffusion constant, D=0 (D), D=0.00005 (E) and D=0.0001 (F). (G) The logarithm of phase diffusion and the logarithm of diffusion coefficient D. The line is a fitting correlation line, and the shaded part is the 95% fitting confidence interval. (H) The change of coherence time and phase coherence when diffusion coefficients (fluctuations) are changed. (I) Sketch map for the definition of phase coherence.

**Figure S11.**
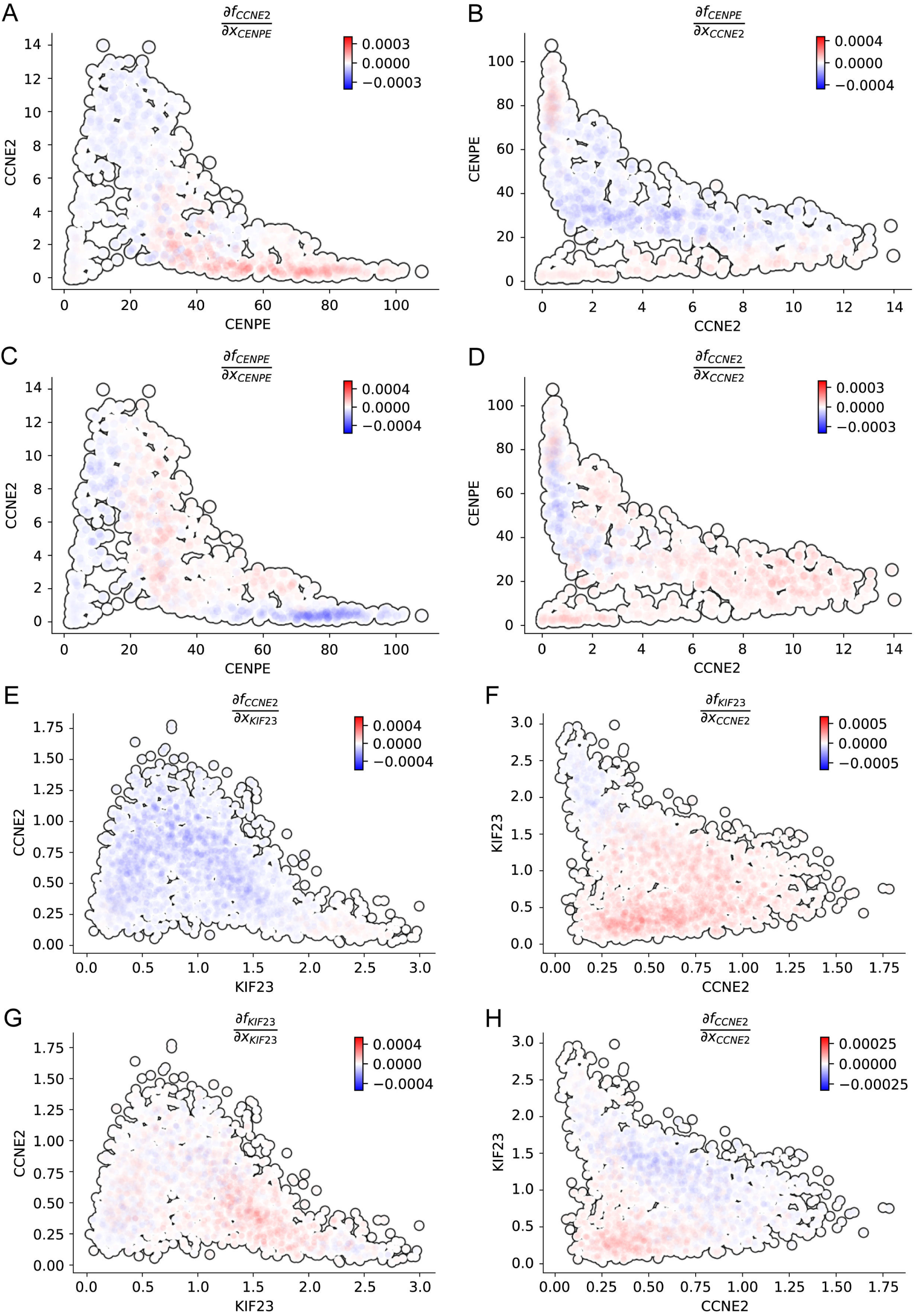
Jacobian analyses of the cell cycle regulatory interactions in the gene expression space (Related to Figure 6) (A-D) Jacobian analyses of the U2OS cell cycle regulatory interactions in the CCNE2 and CENPE expression space. Activation from *CENPE* to *CCNE2* in the *CENPE* and *CCNE2* expression space (A), repression from *CCNE2* to *CENPE* in the *CCNE2* and *CENPE* expression space (B), (C) self-activation of *CENPE* in the *CENPE* and *CCNE2* expression space (C), and self-activation of *CCNE2* in the *CCNE2* and *CENPE* expression space (D). (E-H) Jacobian analyses of the U2OS cell cycle regulatory interactions in the CCNE2 and KIF23 expression space. Repression from *KIF23* to *CCNE2* in the *KIF23* and *CCNE2* expression space (E), activation from *CCNE2* to *KIF23* in the *CCNE2* and *KIF23* expression space (F), self-activation of *KIF23* in the *KIF23* and *CCNE2* expression space (G), and self-activation of *CCNE2* in the *CCNE2* and *KIF23* expression space (H).

**Figure S12.**
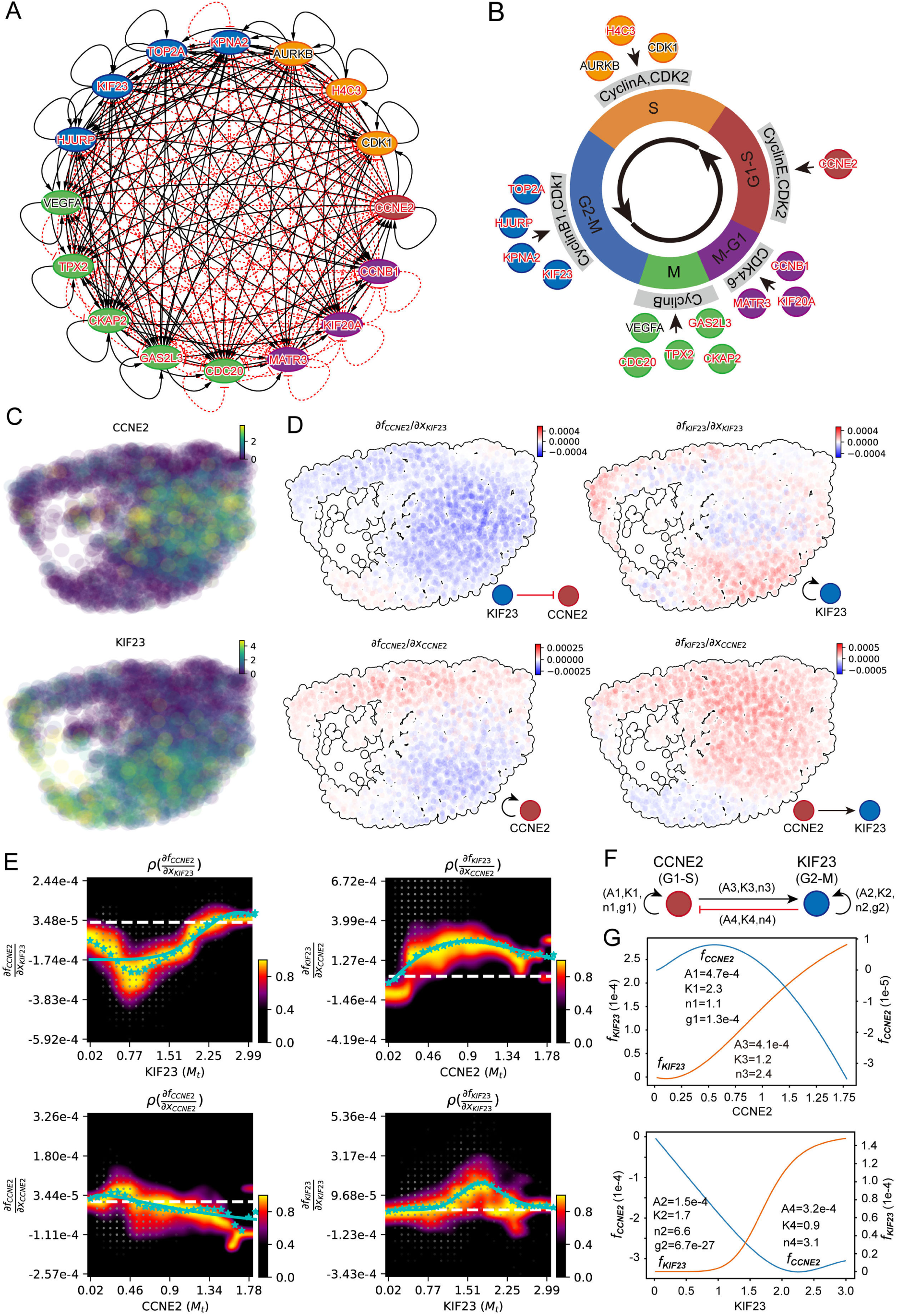
Inference of gene regulatory networks for the RPE1 cell cycle by single-cell transcriptomics data (Related to Figure 6) (A) The interaction network of genes of each cell cycle phase of RPE1 cells. All differentially expressed genes in each cell cycle phase are used to construct network; nodes (genes) with more than one edge are shown. Node colors represent different cycle phases (red: G1-S, yellow: S, blue: G2-M, green: M, purple: M-G1), the black arrows represent the activation and the red arrows represent the inhibition. Genes with black representing TF and red representing non-TFs. (B) The diagram for the cell cycle model with key genes of each cell cycle phase. (C) *CCNE2* has high expression in G1-S phase and *KIF23* has high expression in G2-M phase. (D) Molecular mechanisms underlying the maintenance of the cell cycle. (i) Repression of *CCNE2* by *KIF23*. (ii) Self-activation of *KIF23*. (iii) Self-activation of *CCNE2*. (iv) *CCNE2* activates *KIF23*. (E) Fitting the function of Jacobian versus gene expression with derivatives of a simplistic inhibitory or activation Hill equation. White dashed line corresponds to the zero Jacobian value. The blue stars at each x axis grid point correspond to the weighted mean of the Jacobian values for that point. The blue solid lines are the resultant fittings for the Jacobian. (F) Schematic summarizing the interactions involving *CCNE2* and *KIF23*. (G) The velocity kinetic curves over gene expression changes of the corresponding fitted Hill equations of (E)

